# Role of the *Escherichia coli* ubiquinone-synthesizing UbiUVT pathway in adaptation to changing respiratory conditions

**DOI:** 10.1101/2023.03.15.532739

**Authors:** Arias-Cartin Rodrigo, Kazemzadeh Ferizhendi Katayoun, Séchet Emmanuel, Pelosi Ludovic, Loeuillet Corinne, Pierrel Fabien, Barras Frédéric, Bouveret Emmanuelle

**Affiliations:** Institut Pasteur, Département de Microbiologie, Université Paris-Cité, UMR CNRS 6047, SAMe Unit, Institut Pasteur, France; Univ. Grenoble Alpes, CNRS, UMR 5525, VetAgro Sup, Grenoble INP, TIMC, 38000 Grenoble, France; Univ. Grenoble Alpes, INSERM U1209, CNRS UMR 5309, Institute for Advanced Biosciences, Team Genetics Epigenetics and Therapies of Infertility, 38000 Grenoble, France

**Keywords:** quinone, *E. coli*, Fnr, respiration, UbiTUV

## Abstract

Isoprenoid quinones are essential for cellular physiology. They act as electron and proton shuttles in respiratory chains and in various biological processes. *Escherichia coli* and many α, β, and γ proteobacteria possess two types of isoprenoid quinones: ubiquinone (UQ) is mainly used under aerobiosis, while (demethyl)menaquinones ((D)MK) are mostly used under anaerobiosis. Yet, we recently established the existence of an anaerobic O_2_- independent UQ biosynthesis pathway controlled by *ubiT, ubiU,* and *ubiV* genes. Here, we characterize the regulation of *ubiTUV* genes in *E. coli.* We show that the three genes are transcribed as two divergent operons that are both under the control of the O_2_ sensing Fnr transcriptional regulator. Phenotypic analyses using a *menA* mutant devoid of (D)MK revealed that UbiUV-dependent UQ synthesis is essential for nitrate respiration and for uracil biosynthesis under anaerobiosis, while it contributes, though modestly, to bacterial multiplication in the mouse gut. Moreover, we showed by genetic study and ^18^O_2_ labelling that UbiUV contribute to hydroxylation of ubiquinone precursors through a unique O_2_ - independent process. Last, we report a crucial role of *ubiT* in allowing *E. coli* to shift efficiently from anaerobic to aerobic conditions. Overall, this study uncovers a new facet of the strategy used by *E. coli* to adjust its metabolism upon changing O_2_ levels and respiratory conditions. This work links respiratory mechanisms to phenotypic adaptation, a major driver in the capacity of *E. coli* to multiply in gut microbiota, and of facultative anaerobic pathogens to multiply in their host.

**ABSTRACT IMPORTANCE:** Enterobacteria multiplication in the gastrointestinal tract is linked to microaerobic respiration and associated to various inflammatory bowel diseases. Our study focuses on biosynthesis of ubiquinone (UQ), a key player in respiratory chains, under anaerobiosis. The importance of this study stems from the fact that UQ usage was for long considered to be restricted to aerobic conditions. Here we investigated the molecular mechanism allowing UQ synthesis in the absence of O2 and searched for the anaerobic processes that UQ is fueling in such conditions. We found that UQ biosynthesis involves anaerobic hydroxylases, i.e. enzymes able to insert a O atom in the absence of O2. We also found that anaerobically synthesized UQ can be used for respiration on nitrate and synthesis of pyrimidine. Our findings are likely to be applicable to most facultative anaerobes, which count many pathogens (Salmonella, Shigella, Vibrio) and will help in unravelling microbiota dynamics.

## INTRODUCTION

Isoprenoid quinones are widely distributed in the three domains of life and globally act as electron and proton carriers (1). They serve in many processes of bacterial physiology and electron transport chains like photosynthesis, e.g. plastoquinone and phylloquinone, and respiration, e.g. ubiquinone (UQ) and menaquinone (MK) (2). Isoprenoid quinones are composed of a quinone ring and a poly-isoprenoid side chain whose length varies between organisms (for instance UQ_8_ in *Escherichia coli* and UQ_9_ in *Pseudomonas aeruginosa*). Many proteobacteria, such as *E. coli*, produce two main types of quinones: benzoquinones, represented by UQ, and naphthoquinones, such as MK and demethyl-menaquinone (DMK). In respiratory chains, quinones transfer electrons from primary dehydrogenases to terminal reductases. For decades, *E. coli* aerobic and anaerobic respiratory chains were thought to rely on UQ and MK/DMK, respectively. Yet, we have recently discovered a new pathway for UQ biosynthesis under anaerobiosis, opening the way to a more complex and redundant model for bacterial respiratory metabolism (3).

Aerobic UQ biosynthesis pathway includes 9 steps (4) (Figure S1). It begins with the conversion of chorismate to 4-hydroxybenzoate (4HB) by the chorismate lyase UbiC. Then, the phenyl ring of the 4HB precursor undergoes condensation with a 40-carbon long isoprenoid chain in a reaction catalyzed by the UbiA enzyme. Subsequently, a series of modifications on the 4HB ring by two methylases (UbiE and UbiG), a two-component decarboxylase (UbiD, UbiX), and three hydroxylases (UbiI, UbiH and UbiF) generate the final UQ_8_ product. The FAD-monooxygenases UbiI, UbiH, and UbiF use molecular O_2_ for their hydroxylation reaction (5–7). An atypical kinase-like protein called UbiB is also involved in UQ_8_ synthesis, but its exact role remains elusive (8). In addition, two non-enzymatic factors are required, UbiJ and UbiK, which may allow UbiIEFGH enzymes to assemble in a cytoplasmic 1 MDa complex, referred to as the Ubi metabolon (9). Also, UbiJ and UbiK bind lipids, which may help the hydrophobic UQ biosynthesis to proceed inside a hydrophilic environment.

Anaerobic UQ biosynthesis is formed by a subset of the enzymes of the aerobic pathway, namely UbiA, UbiB, UbiC, UbiD, UbiE, UbiG and UbiX, that function with UbiT, UbiU, and UbiV proteins solely required under anaerobiosis (3) (Figure S1). Like its homolog counterpart UbiJ, UbiT contains a SCP2 lipid-binding domain. Strikingly, UbiU and UbiV do not exhibit any sequence similarity or functional relatedness with the hydroxylases UbiI, UbiH, or UbiF. UbiU and UbiV each contain an iron-sulfur ([4Fe-4S]) cluster coordinated by four conserved cysteine residues embedded in the so-called protease U32 domain, and they form a soluble UbiUV complex (3). Interestingly, two other members of the U32 protein family, RlhA and TrhP, are involved in hydroxylation reactions. They introduce specific nucleotide modifications respectively in the 23S rRNA or in some tRNAs (10–12).

In this work, we aimed at identifying the conditions under which UbiUVT proteins are produced and the genetic regulatory mechanisms involved, and the physiological role of UbiTUV. We concluded that (i) thanks to Fnr control, UbiUV ensure the production of UQ under a range of O_2_ levels, from anaerobiosis to microaerobiosis, (ii) a dual anaerobic/aerobic regulation allows UbiT to secure a rapid shift from anaerobic UbiUV- dependent UQ synthesis to an aerobic UbiIHF-dependent UQ synthesis, and (iii) UbiUV- synthesized UQ can be used for nitrate respiration and anaerobic pyrimidine biosynthesis. We also showed that UbiUV act as O_2_-independent hydroxylases paving the way for future studies towards the characterization of a new type of chemistry.

## RESULTS

### 1. Biochemical function of UbiUV *in vivo*

To get further insight into the UbiUVT system *in vivo*, we tested whether overproduction of UbiU and UbiV could substitute for the three oxygen-dependent hydroxylases UbiI, UbiH, or UbiF. Thus, we cloned the *ubiUV* operon in the pBAD24 vector downstream the arabinose inducible pBAD promoter (pES154 plasmid). In parallel, we also cloned *ubiUV* upstream the SPA tag encoding sequence in order to assess the quantities of proteins produced. The pBAD-*ubiUV*-SPA plasmid produces a level of UbiV protein approximately 30-fold higher to that produced by a chromosomal copy of *ubiV-SPA* under anaerobiosis (Figure S2). After transformation of mutant strains, selection, and precultures with LB medium in absence of O_2_, growth on M9 succinate was tested as it strictly depends upon an aerobic UQ-dependent respiratory chain (Figure 1). In the presence of inducer, the pES154 plasmid was able to suppress the growth phenotype of the Δ*ubiF, ΔubiH, ΔubiIK,* and *ΔubiIHF* mutants (Figure 1A). Note that as a control, we used the *Neisseria meningitidis ubiM* gene that we previously showed to substitute for the growth phenotype of a Δ*ubiIHF* mutant (13). Also, in M9 succinate, the Δ*ubiI* mutation alone has no growth phenotype and needs to be combined with Δ*ubiK* mutation for a defect to be observed (14). To test the importance of the UbiU- bound [Fe-S] cluster, a complementation test was carried out in the same conditions, using a pBAD derivative carrying the *ubiU*(C176A) allele that produces an UbiU variant lacking its [Fe-S] cluster (3). Accordingly, suppression of Δ*ubiH*, Δ*ubiF*, Δ*ubiIK*, and Δ*ubiIHF* was no longer observed (Figure 1A). In addition, the pES154 plasmid was unable to suppress the growth phenotype of Δ*ubiA*, Δ*ubiD*, Δ*ubiE,* or Δ*ubiG* strains (data not shown), and was also unable to suppress the growth phenotype of Δ*ubiH*Δ*ubiA* or Δ*ubiH*Δ*ubiD* mutants (Figure 1B), showing that UbiUV intervene specifically at the hydroxylation steps and otherwise depend upon all the other components of the aerobic UQ biosynthesis pathway to do so. These results indicate that in the presence of O_2_, expression of UbiUV can substitute for the O_2_-dependent UbiIHF hydroxylases and that integrity of the UbiU [Fe-S] cluster is required.

**Figure 1.**
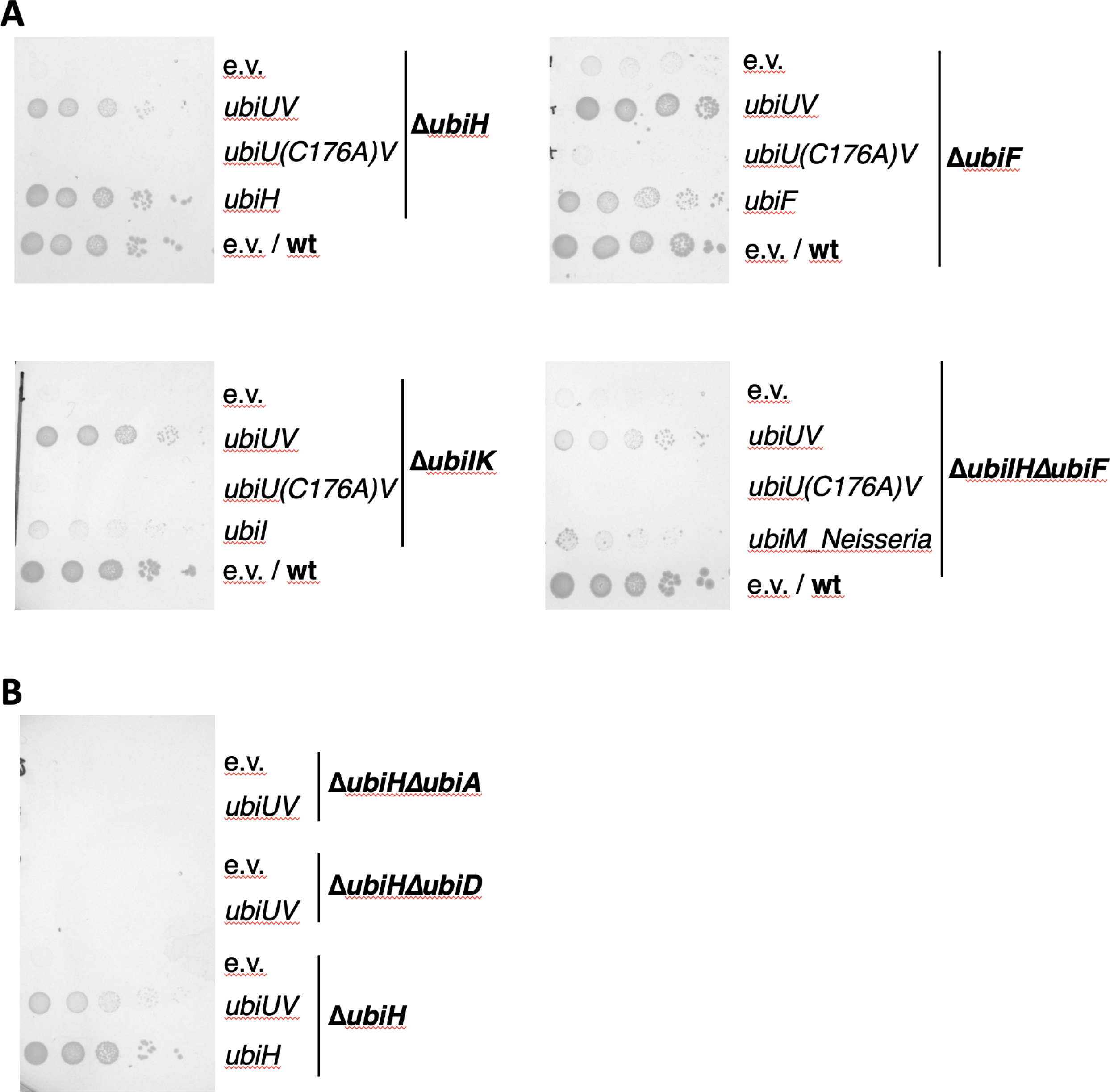
Complementation of *ubiI, ubiH,* and *ubiF* mutants by pBAD-*ubiUV* in the presence of O2. (A). *E. coli* mutant strains Δ*ubiH* (FBE253), Δ*ubiF* (FBE512), Δ*ubiIK* (FBE713), and Δ*ubiIHΔubiF* (FBE650) were transformed by pBAD24 (e.v.), pBAD-UbiUV, and pBAD- UbiU(C176A)V plasmids. (**B).** *E. coli* mutant strains Δ*ubiHΔubiA* (FBE792), Δ*ubiHΔubiD* (FBE793), and Δ*ubiH* (FBE253) were transformed by pBAD24 (e.v.) and pBAD-UbiUV. (A-B) After selection in the absence of O_2_, cultures were washed and serially diluted in minimal medium then spotted on M9 succinate plates containing 0.02% arabinose, and incubated at 37°C for 48 hours (or 96 hours for the Δ*ubiIHF* series) in aerobic conditions (21% O_2_). The results shown are representative of at least two independent experiments.

Remarkably, expression of the pES154 plasmid was also able to suppress growth defects of the Δ*ubiJ* mutant (Figure 2A). UbiJ is an auxiliary factor important for organizing the aerobic Ubi metabolon. We reasoned that suppression was made possible thanks to the presence of the chromosomally encoded UbiT that shares sequence similarity with UbiJ. To test this, we repeated the complementation test in two new strains, Δ*ubiH*Δ*ubiJ* and Δ*ubiH*Δ*ubiT*. The pES154 plasmid still complemented growth defects of the Δ*ubiH*Δ*ubiJ* mutant, but it was unable to complement the Δ*ubiH*Δ*ubiT* mutant (Figure 2B). Similarly, pES154 was found to suppress the growth defect phenotype of a Δ*ubiF*Δ*ubiJ* mutant but not a Δ*ubiF*Δ*ubiT* mutant (Figure 2C). These results showed that in the presence of O_2,_ increased dosage of *ubiUV* genes suppresses the lack of O_2_-dependent hydroxylases UbiF and UbiH in an UbiT-dependent/UbiJ-independent manner.

**Figure 2.**
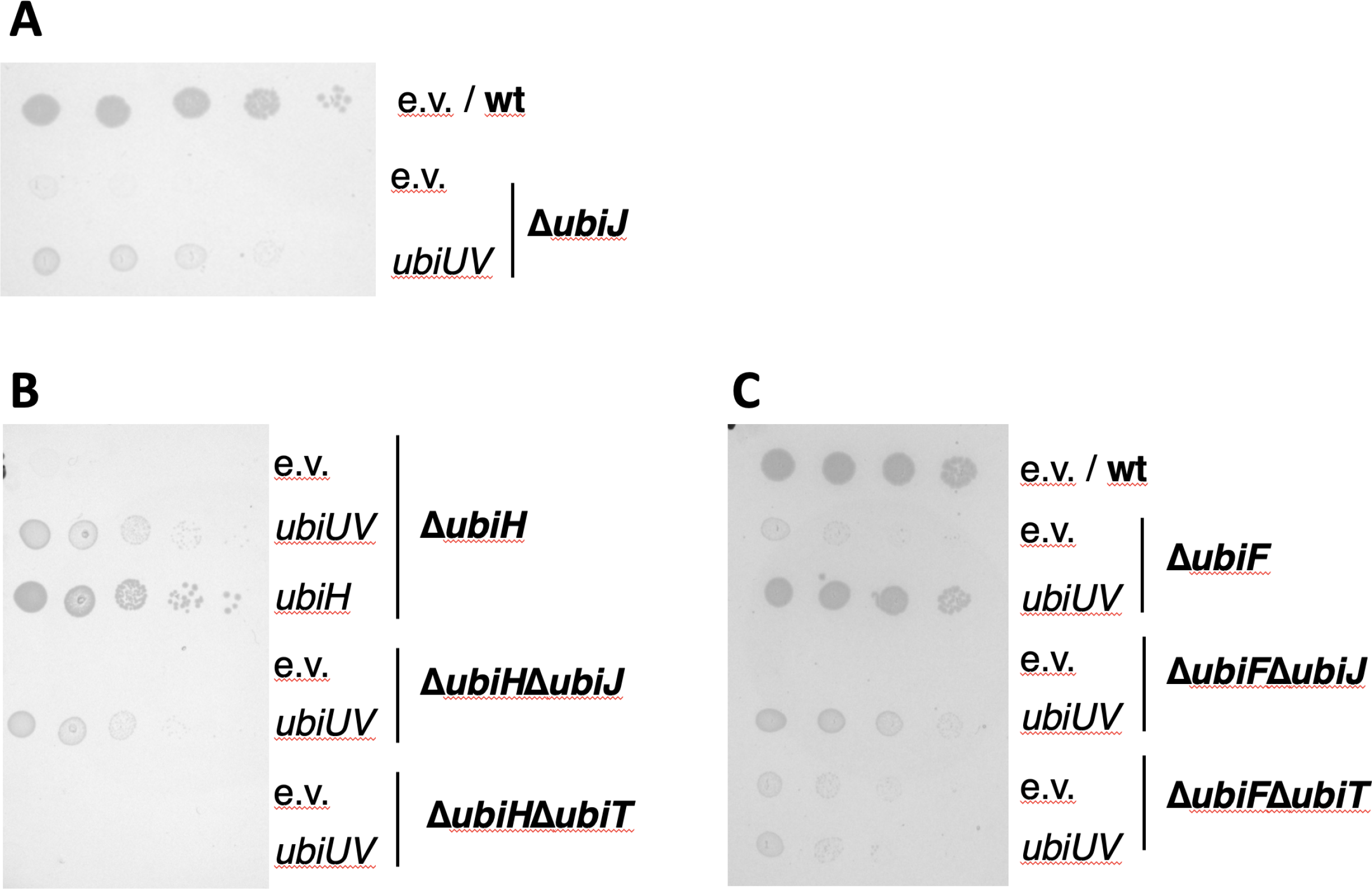
Complementation of *ubiH* and *ubiF* mutants by pBAD-UbiUV is *ubiT* dependent (and *ubiJ* independent). *E. coli* mutant strains were transformed by pBAD24 (e.v.) and pBAD- UbiUV plasmids. After selection in the absence of O_2_, cultures were washed and serially diluted in minimal medium then spotted on M9 succinate plates containing 0.02% arabinose at 37°C in aerobic conditions. The results shown are representative of at least two independent experiments. **(A)** strains Δ*ubiJ* (FBE514) and wt **(B)** strains Δ*ubiH* (FBE253), Δ*ubiHΔubiJ* (FBE794), and Δ*ubiHΔubiT* (FBE795); **(C)** strains Δ*ubiF* (FBE512), Δ*ubiFΔubiJ* (FBE264), Δ*ubiFΔubiT* (FBE265), and wt.

To confirm that phenotypic suppression was due to UQ_8_ synthesis, we quantified the UQ_8_ content by HPLC analysis coupled to electrochemical detection (ECD) for all strains described above (Figure 3A). Results showed that mutant strains lacking UbiI-UbiK, UbiH and/or UbiF were severely deficient in UQ. The pES154 plasmid enabled Δ*ubiH*, Δ*ubiF,* or Δ*ubiIH* strains to synthesize 30-50% of the UQ level of the wild-type (wt) strain (Figure 3A, 1^st^ panel). The levels of UQ obtained in the Δ*ubiIHΔubiF* and Δ*ubiIK* mutant strains with the pES154 plasmid were much lower. We stress that the UQ levels cannot be directly correlated with the phenotypic analysis (Figures 1-2) since culture media were different (LB vs M9 succinate) to allow recovery of enough biological material for the HPLC-ECD analyses. Importantly, the pBAD-*ubiU(C176A)V* plasmid was unable to promote UQ synthesis in Δ*ubiH* (Figure 3A, 2^nd^ panel). Last, UQ_8_ content assay confirmed that UbiT, but not UbiJ, was necessary for UbiUV to synthesize UQ in aerobic conditions (Figure 3A, 3^rd^ panel).

**Figure 3.**
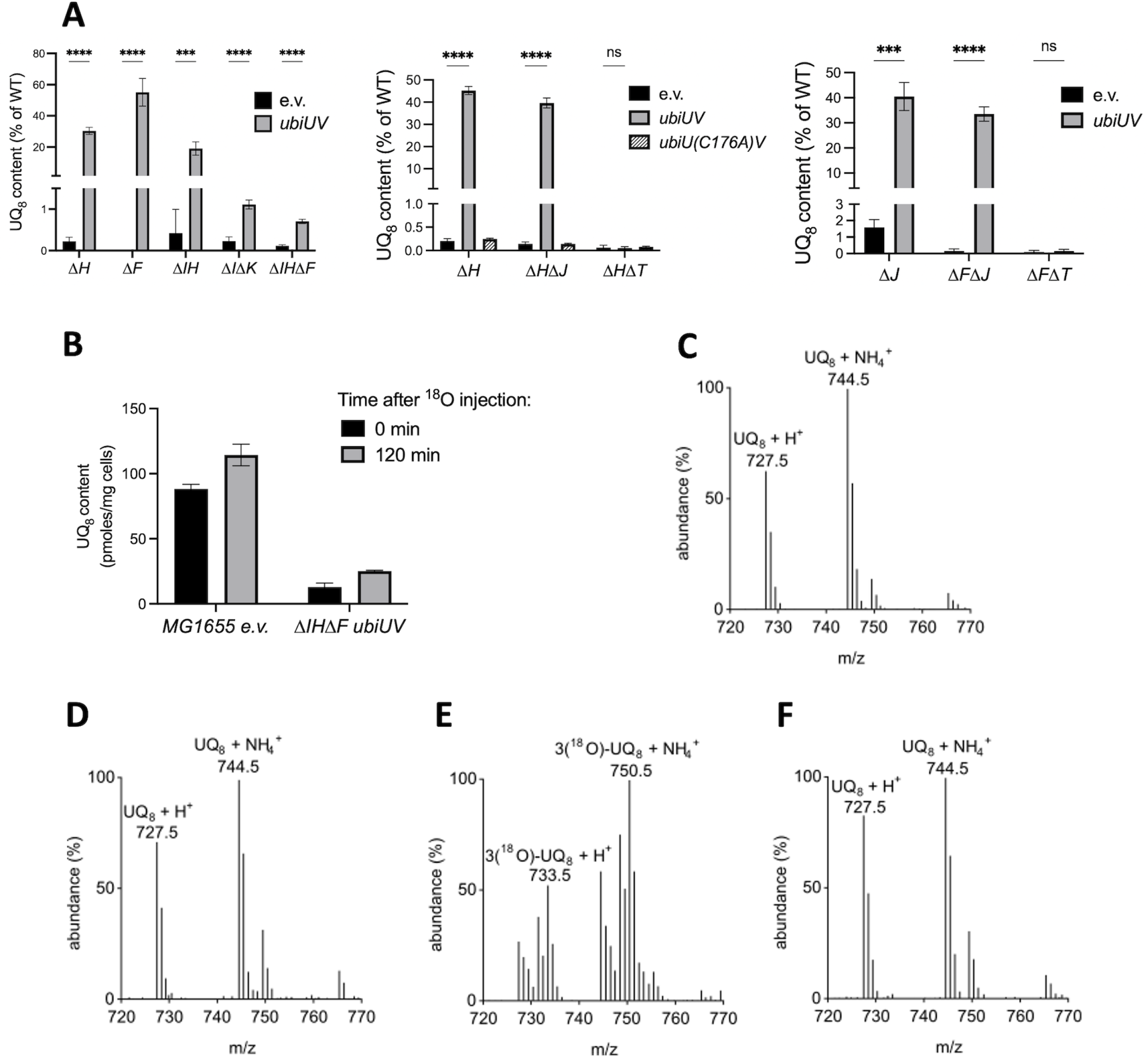
UbiUV restore the UQ_8_ content of Δ*ubiIH* and *ΔubiF* mutants without using O_2_ for the hydroxylation steps. (A) UQ_8_ content of the indicated strains containing either pBAD (e.v.), pBAD-*ubiUV,* or pBAD-*ubiU(C176A)V* after aerobic growth overnight at 37°C in LB medium. *E. coli* wt strain (MG1655) containing the empty vector was used as control. Mean ± standard deviations (SD) (n=3 to 4). ***, P < 0.001; ****, P < 0.0001 by unpaired Student’s t test. **(B-F)** Detection of UQ_8_ with ^18^O_2_ labelling. **(B)** Quantification of UQ_8_ content in wt (MG1655) cells containing an empty vector (e.v.) and in Δ*ubiIHΔubiF cells* containing the pBAD-*ubiUV* vector just before (0 min) and two hours (120 min) after adding ^18^O_2_. Mean ± SD (n=2). (**C to F**) Mass spectra of UQ_8_ from cells shown in B, wt (**C and E**) and Δ*ubiIHΔubiF* with pBAD-*ubiUV* (**D and F**), before (**E and F**) and two hours after (**E and F**) addition of ^18^O_2_. Mass spectra representative of two independent experiments.

The results above showed that UbiUV hydroxylate UQ precursors, when expressed under aerobic conditions. This result raised the possibility that under such conditions, O_2_ might be used as a co-substrate of the hydroxylation reactions, as is the case for UbiI, UbiH and UbiF in wt cells (5). To test this hypothesis, we exposed cells to ^18^O_2_ and monitored the labelling of UQ by HPLC-ECD-MS. Two hours after ^18^O_2_ addition, the level of UQ_8_ increased in both strains (Figure 3B). Before adding ^18^O_2_, the mass spectra of UQ synthesized by wt or Δ*ubiIH*Δ*ubiF* cells containing pES154 displayed H^+^ and NH_4_^+^ adducts with m/z ratio characteristic of unlabeled UQ (Figure 3C and D). As expected, two hours after adding ^18^O_2_, most of the UQ_8_ pool in wt cells contained three ^18^O_2_ atoms (Figure 3E), in agreement with O_2_ being the co-substrate of the aerobic hydroxylation steps (5). In contrast, we detected only unlabeled UQ_8_ in the Δ*ubiIH*Δ*ubiF* strain expressing UbiUV (Figure 3F), demonstrating that UbiUV utilize another oxygen donor than O_2_, even when operating under aerobic conditions.

Altogether, both phenotypic and UQ_8_ quantification results allowed us to conclude that UbiU and UbiV, when produced in sufficiently high-level, function in the canonical “aerobic” UQ_8_ biosynthesis pathway by catalyzing [Fe-S]-dependent hydroxylation of the benzene ring in an O_2_-independent reaction. Remarkably, UbiT is necessary for such aerobic UbiUV-mediated synthesis to occur and cannot be substituted by UbiJ.

### 2. The ISC [Fe-S] biogenesis machinery is required for anaerobic UQ biosynthesis

The UbiU and UbiV proteins each contain a [4Fe-4S] cluster, which is essential to the synthesis of UQ in anaerobic conditions (3). Assembly of [4Fe-4S] clusters requires complex biosynthetic machineries, ISC and SUF (15). Therefore, the UQ_8_ levels were monitored in Δ*isc* and Δ*suf* mutants grown in anaerobic conditions (Figure S3). In addition to UQ_8_, the quantification of DMK_8_ and MK_8_ was also performed. UQ_8_ content in Δ*isc* mutants was strongly impaired (around 15% of the wt), and was much less affected in Δ*suf* mutants (60- 80% of the wt). DMK and MK content remained mostly unaltered. Collectively, these results showed that ISC and SUF systems are not involved in DMK and MK biosynthesis and that the ISC system is the most relevant in UQ_8_ biosynthesis likely through the maturation of [4Fe-4S] clusters in UbiU and UbiV.

### 3. Anaerobic and micro-aerobic UQ biosynthesis

Genome scale studies have predicted that *ubiUV* genes are under the control of the anaerobic Fnr transcriptional activator (16, 17). In contrast, *ubiT* did not appear as a potential Fnr target. This prompted us to investigate the effect of anaerobiosis (0% O_2_), microaerobiosis (0.1% O_2_), and aerobiosis (21% O_2_) on the level of UbiU, UbiV, and UbiT proteins. In order to follow the quantity of UbiTUV proteins in physiological conditions, we constructed a series of recombinant strains producing the UbiT, UbiU, or UbiV proteins with a C-terminal SPA tag (18) encoded from a gene fusion at their chromosomal loci. We examined protein production by Western Blot assay using an anti-flag antibody and assessed loading with a polyclonal antibody against YbgF (CpoB). All three UbiTUV-SPA tagged proteins were present in strains grown in anaerobiosis (Figure 4A) and microaerobiosis (Figure 4B). In aerobiosis, production of UbiU and UbiV was no longer observed whereas a significant level of UbiT was still visible. The contribution of Fnr to anaerobiosis- or microaerobiosis-mediated activation of *ubiU* and *ubiV* genes was confirmed as no cognate UbiU or UbiV associated band was observed in a Δ*fnr* mutant (Figure 4A-B). Interestingly, UbiT level was also reduced in the Δ*fnr* mutant in -O_2_. Last, in order to validate the physiological significance of the Fnr regulatory circuit depicted above, we quantified the amount of UQ_8_ produced in wt and Δ*fnr* strains, during aerobiosis and anaerobiosis (Figure 4C). In comparison to the UQ content found in the wt strain in aerobiosis, the level in anaerobiosis was reduced by half. Importantly, we observed that almost no UQ was detected in the Δ*fnr* mutant (Figure 4C). This revealed the pivotal role that Fnr plays in allowing UQ_8_ synthesis in the absence of O_2_.

**Figure 4.**
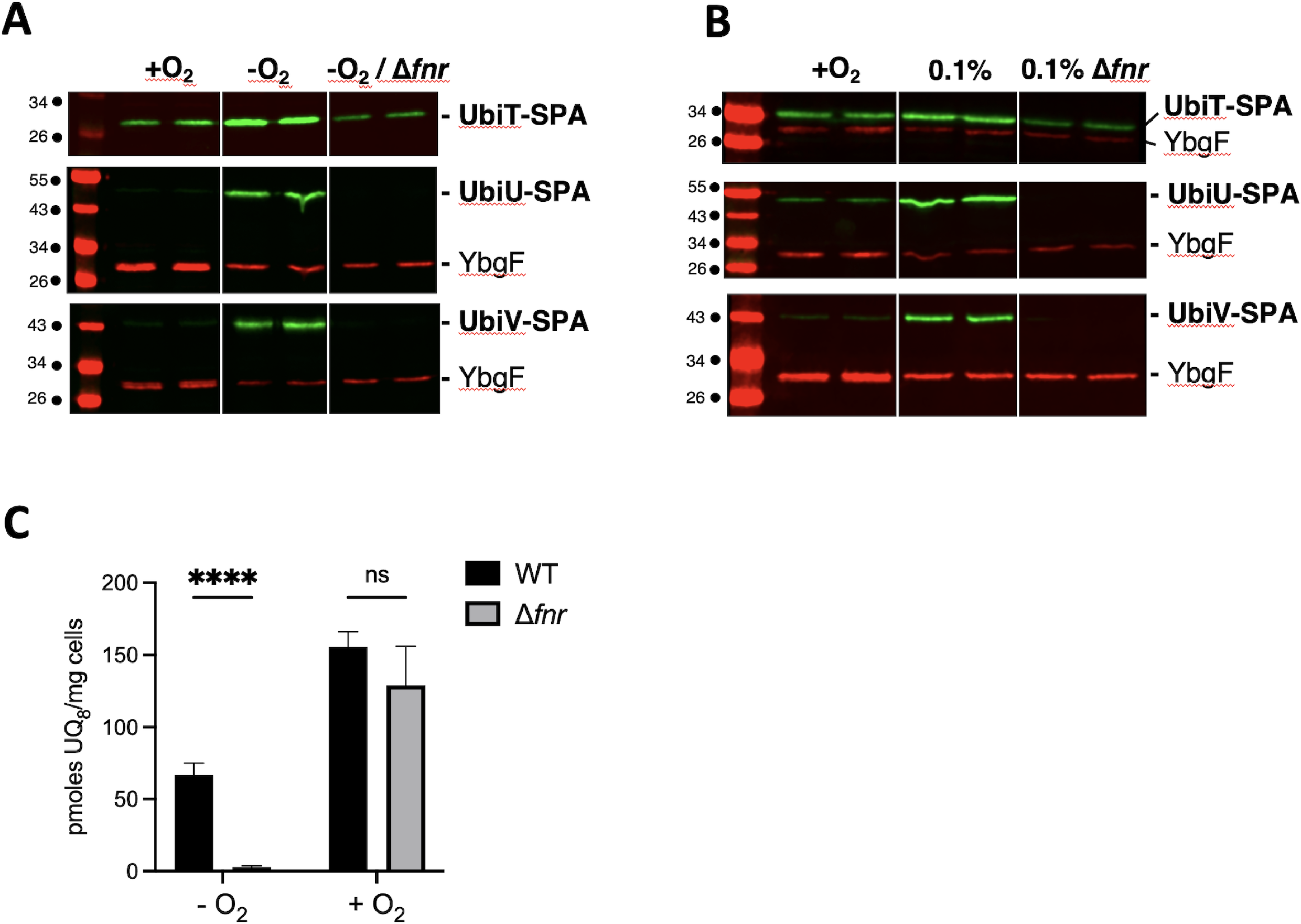
Fnr controls UbiTUV expression and UQ biosynthesis under anaerobiosis. (A.B). *E. coli* strains UbiU-SPA, UbiV-SPA, and UbiT-SPA, and their corresponding Δ*fnr* versions (strains FBE656, FBE789, FBE655, FBE695, FBE696, FBE694) were grown in LB at 37°C in +O_2_ and –O_2_ (**A**) or in +O_2_ and 0.1% O_2_ (**B**) and analyzed by Western blot: normalized quantities of total protein extracts (in biological duplicate) were separated by SDS-PAGE 12% and detected by Western-Blot using anti-Flag monoclonal antibody for the detection of the SPA tag (green) or anti-YbgF polyclonal antibodies as an internal loading control (red). **(C)**. UQ_8_ content of the wild type and Δ*fnr* (FBE354) strains was assayed after aerobic or anaerobic growth overnight at 37°C in LB medium. Mean ± standard deviations (SD) (n=3 to 4). ****, P < 0.0001 by unpaired Student’s t test.

### 4. Genetic control of *ubiUVT* gene expression

Previous genome Chip-seq analysis reported binding of Fnr within the *ubiT*-*ubiUV* intergenic region. Additionally, in a whole genome sequence search study, one transcription start site has been described upstream of the *ubiUV* operon (*ubiUV_p_*) and two sites described upstream of *ubiT* (*ubiT_p1_*, *ubiT_p2_*) (19) (Figure 5A, 5B). Upon inspection of that region, we were able to identify two potential Fnr binding sites fitting well with the described Fnr binding consensus. The F1 site, reading TTGATTTAAGGCAG is located 36 nucleotides (nt) upstream the *ubiUV_p_*transcription start site (Figure 5A). The F2 site, reading TTGATTTATACCGC locates 33 nt upstream the proximal +1 transcription starting site *ubiT_p2_* and 19 nt downstream the distal *ubiT_p1_*(Figure 5A, 5B).

**Figure 5.**
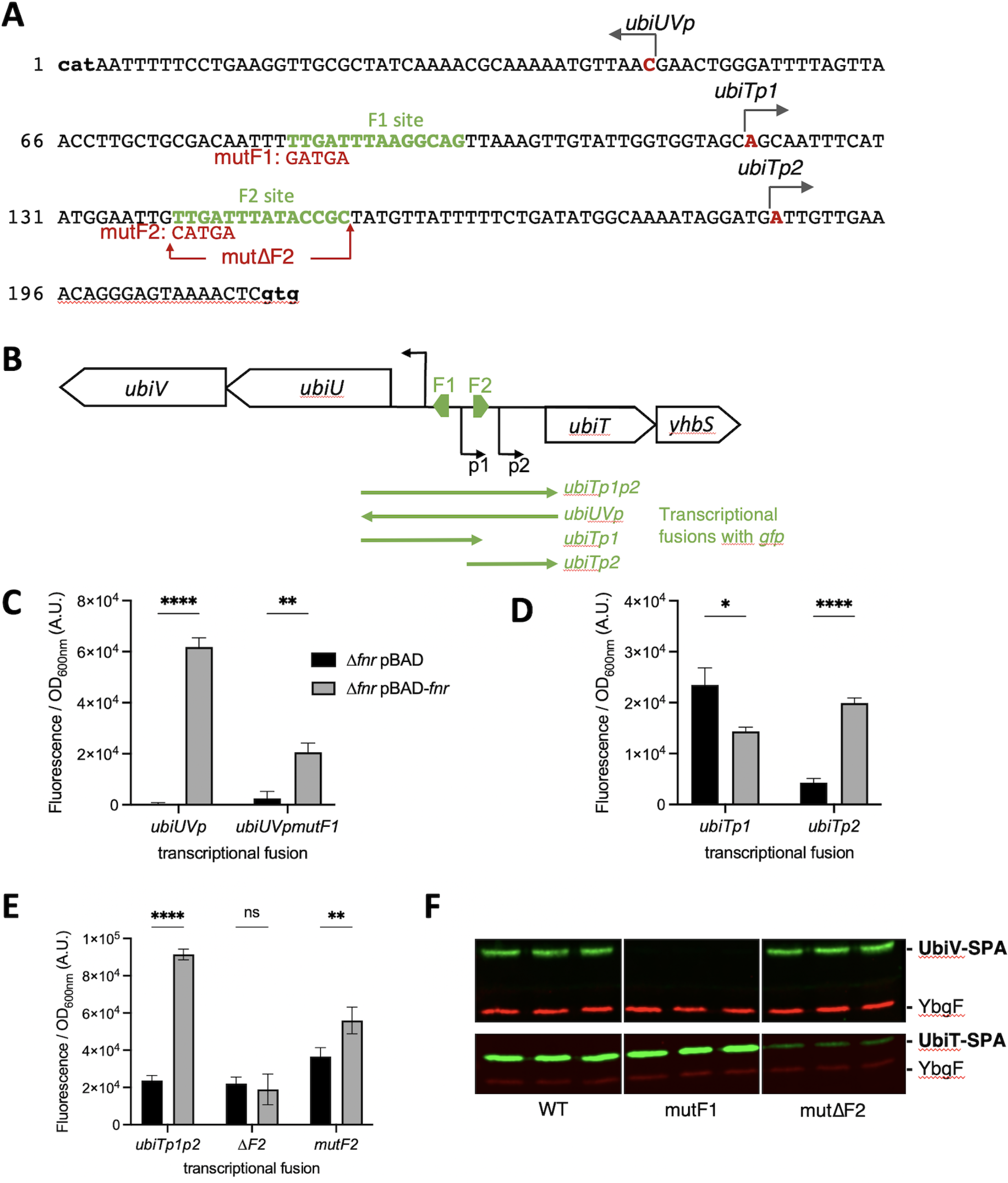
(A). Organization of the promoter region of *ubiTUV* genes. The sequence of the intergenic region between *ubiU* and *ubiT* genes is shown, from the start codon of *ubiU*, to the start codon of *ubiT* (both indicated in bold at the extremities of the sequence). The transcription start sites as determined in (19), are indicated in red. The two Fnr binding sites F1 and F2 are indicated in green. The mutF1, mutF2, and mutΔF2 mutations introduced in the transcriptional fusions are depicted in red. (**B). Limits of transcriptional fusions used in panels C-E. (C-E). Activity of the transcriptional fusions with or without *fnr* expression.** Δ*fnr E. coli* strain was co-transformed by pBAD24 or pBAD-fnr together with the indicated transcriptional fusions. After overnight growth of 4 biological replicates et 37°C in LB in anaerobiosis, GFP levels were quantified. Errors bars indicate the standard deviation. *, P < 0.1; **, P < 0.01; ****, P < 0.0001 by unpaired Student’s t test. (**F). Role of the Fnr sites in UbiTUV physiological levels.** *E. coli* strains UbiV-SPA and UbiT-SPA, without (wt) or with the indicated mutation in the F1 or F2 binding sites, were grown in LB overnight at 37°C in the absence of O_2_. Normalized quantities of total protein extracts (in biological triplicate) were separated by 12% SDS-PAGE and detected by Western-Blot using anti-Flag monoclonal antibody for the detection of the SPA tag (green) or anti-YbgF polyclonal antibody as an internal loading control (red).

To detail the molecular mechanism of regulation and to dissect the promoter organization of the intergenic region between *ubiUV* and *ubiT*, we used transcriptional fusions with GFP (20). We used 4 different transcriptional fusions encompassing *ubiUV_p_, ubiT_p1_, ubiT_p2_*, and a construction *ubiT_p1p2_*containing the two promoters of *ubiT* (Figure 5B). We compared the expression of these transcriptional fusions in anaerobiosis, in a Δ*fnr* mutant complemented or not with a pBAD-*fnr* plasmid. The *ubiUV_p_* and *ubiT_p2_* promoters were strongly activated in the presence of pBAD-*fnr*, whereas the *ubiT_p1_*promoter was not (Figure 5C-D). This suggested that the *ubiUV_p_* promoter was activated by Fnr binding to the F1 site, and that the *ubiT_p2_*promoter was activated by Fnr binding to the F2 site. When we introduced mutations in the F1 binding site (5 mutated nucleotides; mutF1; Figure 5A), the activation of the expression from the *ubiUVp* transcriptional fusion was severely reduced (Figure 5C). Mutations of the F2 site (mutΔF2 complete deletion or mutF2 with 5 mutated nucleotides; Figure 5A) also affected the expression of the ubiT*_p1p2_* transcriptional fusion, but a basal level of expression was maintained, probably due to the expression from the distal *ubiT_p1_* promoter (Figure 5E).

Next, we introduced the same mutations in the F1 and F2 Fnr binding sites at the locus in the *ubiU-ubiT* intergenic region in the chromosome of the strains producing UbiV- SPA or UbiT-SPA tagged proteins. Mutation within the F1 site upstream *ubiU* completely prevented the production of UbiV in the absence of O_2_ (Figure 5F). Mutation within the F2 site upstream the proximal *ubiT_p2_* promoter prevented the induction of *ubiT* in the absence of O_2_, without altering the basal level of UbiT-SPA observed in the presence of O_2_ (Figure 5F and Figure S4). Notably, the mutation in the F1 binding site did not affect the expression of *ubiT* and conversely, mutation of the F2 binding site did not affect the expression of *ubiV*.

Altogether, these results showed that Fnr activates *ubiUV* transcription under anaerobiosis, while *ubiT* expression can be triggered from two promoters, one aerobically active (P1) and the other anaerobically active (P2) under Fnr control.

### 5. Physiological role of UbiUVT at different O_2_ levels

We have previously reported that UbiU, UbiV, and UbiT are essential for anaerobic synthesis of UQ in *E. coli* when grown in LB, glycerol/DMSO, or lactate/NO_3_^-^ (3). However, the contribution to *E. coli* physiology of UQ synthesized by UbiUVT in anaerobic conditions was not investigated in detail. We made use of a set of mutants altered in aerobic (*ubiH*) or anaerobic (*ubiUV*, *ubiT*) UQ_8_ synthesis, as well as mutants altered in DMK/MK biosynthesis (*menA*) to assess the contribution of each type of quinone for growth in a wide range of O_2_ level, 21% (aerobic), 0.1% (microaerobic), and 0% O_2_ (anaerobic), and with varying carbon sources (e.g. glycerol or glucose) and electron terminal acceptors (e.g. O_2_ or NO_3_^-^).

In the presence of glycerol and NO_3_^-^ under aerobic conditions (Figure 6, upper left panel) Δ*ubiUV* and Δ*ubiT* strains showed no growth phenotype. In such conditions, while NO_3_ is present, O_2_ is used for respiration. This contrasted with the Δ*ubiH* mutant, which was severely affected. Combining Δ*ubiH* and Δ*ubiUV* bore no aggravating effect. In contrast, combining both Δ*ubiH* and Δ*menA* had an aggravating effect, indicating that in addition to UQ, DMK and/or MK can support *E. coli* growth even in aerobiosis, as previously suggested (21). In microaerobic conditions (Figure 6, upper center panel), no phenotype was observed for Δ*ubiUV* or Δ*ubiT* strains. In contrast, the Δ*menA* Δ*ubiH* strain still exhibited a clear defect, suggesting that *ubiUV* and *ubiT* do not bear a prominent role in NO_3_^-^-dependent respiratory metabolism under microaerobic conditions, despite being expressed in microaerobiosis (see above). This notion was also supported by the fact that at 0.1% O_2,_ the Δ*ubiH* and Δ*ubiH* Δ*ubiUV* strains did not show any phenotype. At 0.1% O_2_, UQ-dependent metabolism through cytochrome *bd* or *bo* oxidases would remain inconsequential and cells presumably rely on DMK/MK-dependent metabolism for anaerobic respiration (22). Last, in anaerobic conditions, with NO_3_^-^ used for respiration, Δ*ubiUV*, Δ*ubiT* and Δ*menA* strains showed wt-like growth phenotype (Figure 6, upper right panel). However, combining Δ*menA* and Δ*ubiUV* mutations or Δ*menA* and Δ*ubiT* mutations drastically hampered NO_3_^-^ respiratory capacities. In fact, growth of these mutants on M9 glycerol NO_3_^-^ was barely better than a Δ*fnr* strain (Figure 6), which was used as control since it was shown that such strain is unable to respire nitrate but can still use glucose anaerobically (23). These results indicated that anaerobically UbiUVT-synthesized UQ and MK are fully interchangeable electron carriers during NO_3_^-^ respiration under full anaerobiosis (24). Furthermore, we could exclude that the aerobic UQ biosynthetic pathway could contribute to growth in such conditions as the Δ*ubiH* and Δ*menA* Δ*ubiH* mutants exhibited no growth phenotype.

**Figure 6.**
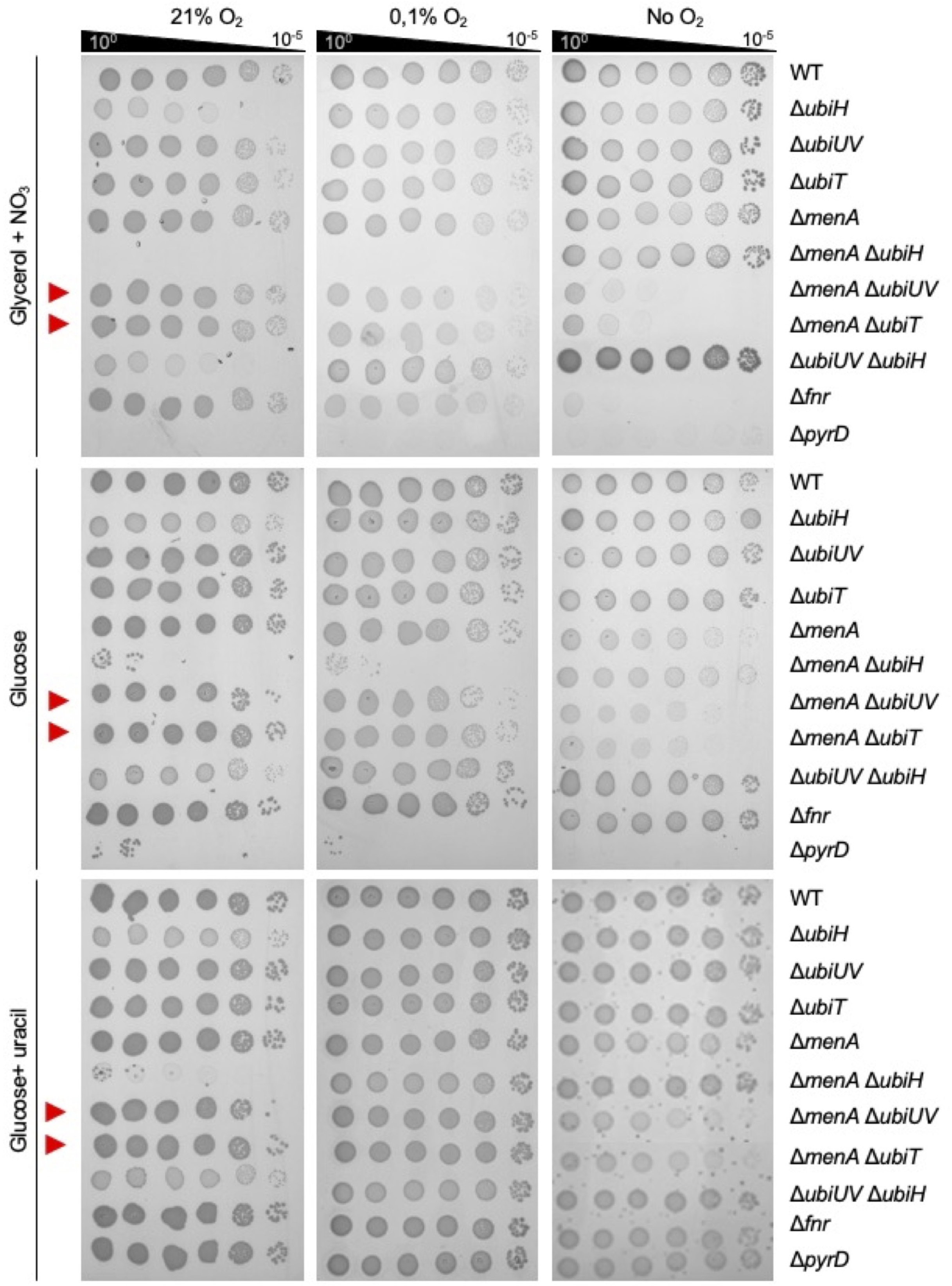
Role of *ubiUVT* in anaerobic growth. *E. coli* wt and strains devoid of the MK/DMK (Δ*menA),* UQ synthesis pathways – aerobic (Δ*ubiH*) and anaerobic (Δ*ubiUV* or Δ*ubiT*) – and controls for anaerobic growth (Δ*fnr*) and auxotrophy for uracil (Δ*pyrD*) were grown aerobically at 37°C in LB medium or LB glucose 0.2% (for Δ*menA ΔubiH*), washed and resupended in M9 medium without carbon source to OD_600_ 1. Serial dilutions were spotted in agarose M9 medium plates supplemented with carbon source (glycerol or glucose), KNO_3_ or Uracil and incubated at 37°C at the indicated O_2_ concentration until growth was observed. Experiments were performed in triplicates and confirmed with at least 4 independent biological replicates.

In the presence of glucose as a carbon source and under aerobiosis, Δ*ubiUV* and Δ*ubiT* mutants exhibited wt-like growth capacity (Figure 6, left middle panel). The Δ*ubiH* mutant showed some slower growth but a most spectacular negative additive effect was observed upon combining Δ*ubiH* and Δ*menA* mutations. It likely points out a role for DMK/MK in aerobic electron transport (25). In anaerobiosis, neither Δ*ubiH* nor Δ*menA*, alone or in combination, showed defect in the presence of glucose as a carbon source (Figure 6, middle right panel). In contrast, Δ*menA* Δ*ubiUV* or Δ*menA* Δ*ubiT* mutants exhibited additive growth defect (Figure 6, middle right panel). This indicated that the UbiUVT-biosynthesized UQ was crucial for growth in glucose fermentative conditions, in the absence of MK. A possibility was that this negative effect reflected auxotrophy for uracil, whose synthesis depends upon electron transfer from PyrD dihydrooratate dehydrogenase to fumarate reductase (FrdABCD) via quinones in anaerobiosis (26). As a matter of fact, adding uracil to the medium had a rescuing effect (Figure 6, lower right panel), supporting the notion that uracil deficiency was responsible for the growth defect observed in the Δ*ubiUV* Δ*menA* mutant in anaerobiosis. This was an important observation as early studies had proposed that the PyrD/FrdABCD electron transfer chain relied mostly on MK/DMK and marginally, if at all, on UQ (26). Our observation clearly shows that anaerobically synthesized UQ can also allow functioning of PyrD. Incidentally, we noticed that the addition of uracil did not rescue the growth defect of the Δ*menA*Δ*ubiH* mutant in aerobiosis, but we have no explanation for this observation.

### 6. Contribution of the O_2_-independent UQ biosynthesis pathway to mouse intestine colonization

Since enterobacteria evolve mostly in anaerobic conditions in their natural habitat, we evaluated the physiological importance of the O_2_-independent UQ biosynthesis pathway in the mouse intestine. To do so, we performed competition experiments between two isogenic strains, MP7 and MP13, which respectively express mCherry and GFP in the presence of tetracycline (27). We deleted *ubiUV* in the MP13 background and confirmed, as expected, that this strain was deficient for UQ_8_ when grown anaerobically (Figure S5A). MK was previously shown to be important for efficient colonization of the mouse intestine by *E. coli* (28). Thus, we also constructed a Δ*menA* mutant in the MP13 background. We checked that deletion of Δ*menA* abrogated the synthesis of DMK and MK (Figure S5B and C). The fitness of the Δ*ubiUV* and Δ*menA* mutants was tested in competition experiments with the MP7 wt strain. We monitored the abundance of each strain in the feces of mice up to 10 days after co-inoculation by oral gavage (Figure 7A). In both experiments, the total CFU count reached ∼10^8^ per gram of feces 24 hours post inoculation (Figure 7B and C and S6A and B) and then gradually decreased to ∼10^5^, showing efficient colonization of the MP7 strain. The abundance of the *ubiUV* mutant was slightly decreased compared to wt (Figure 7B and S6A), which translated in an average competitive index (CI) <1 (Figure 7D and S6C) at days 1, 2, 4, and 10. We noticed however a rather high inter-individual variability (Figure S6C). In contrast, the Δ*menA* mutant was markedly less abundant than the wt (Figure 7C and S6B) and was even undetectable at day 10. CI <1 were observed for every mouse at every sampling (Figure 7E and S6D) and the values obtained were much lower than in the case of the Δ*ubiUV* mutant. Collectively, these data confirm that DMK/MK is the most important quinone for the physiology of *E. coli* in the mouse intestine (28). However, they also reveal a contribution, albeit minor, of the O_2_-independent UbiUV-mediated UQ biosynthesis pathway.

**Figure 7.**
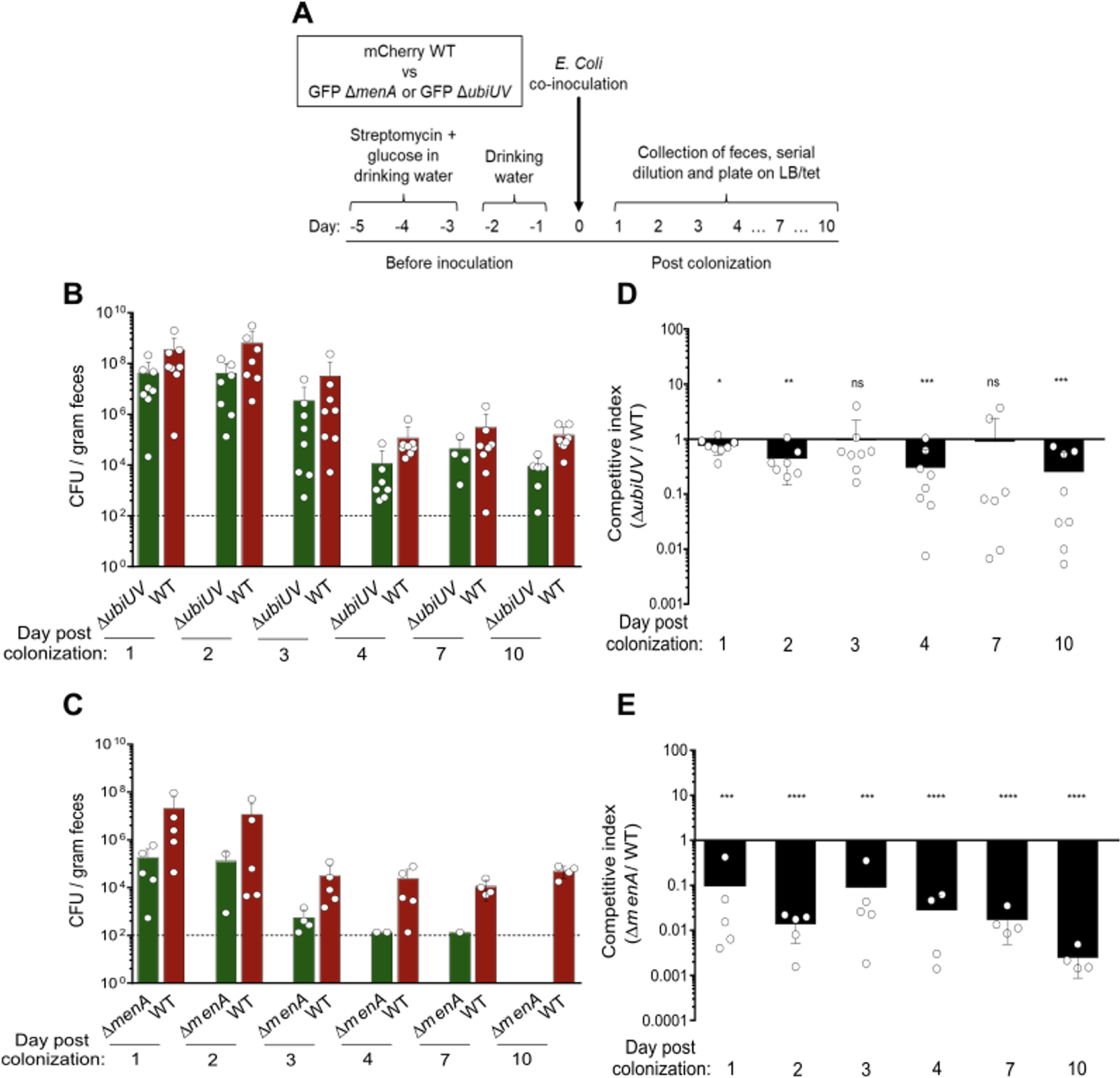
Quinones contribute differently to colonization of the mouse gut by *E. coli*. (A). Schematic representation of the experimental protocol for the mouse intestine colonization competitions, adapted from (42). (**B-C**) Total CFU counts (**B**) and associated competitive index (**C**) in feces of mice after oral co-inoculation with a 1:1 ratio of MP7 wt and MP13 Δ*ubiUV* strains. (**D-E**) same as B-C with MP7 wt and MP13 Δ*menA* strains. The limit of detection of 10^2^ CFU is indicated as dotted line. Mean ± standard deviations (SD), each white circle represents values for individual mice (n=5 and 8), circles missing corresponds to the absence of feces for that day. ns, not significant; *, P < 0.05; **, P < 0.01; ***, P < 0.001; ****, P < 0.0001 by one-sample t test. Changes in total CFU counts and CI throughout the experiment in each mouse are shown in Figure S6.

### 7. Role of UbiT within the anaerobiosis-aerobiosis shift

Phenotypic analysis above revealed that anaerobically UbiUVT-synthesized UQ was contributing to growth via glucose fermentation or NO_3_^-^ respiration. In both conditions, anaerobic UbiUVT-synthesized UQ was functionally redundant with anaerobically synthesized DMK/MK. Because UQ is crucial under aerobiosis, we reasoned that anaerobically synthesized UQ might prepare the cells to adapt to an aerobic environment, i.e. before the aerobic UbiIHF-dependent synthesis takes over. Thus, we investigated the role of UbiUVT-synthesized UQ in the anaerobiosis-aerobiosis transition.

Firstly, we used Δ*menA*Δ*ubiH* and Δ*menA*Δ*ubiUV* strains that only produce UQ under anaerobiosis and aerobiosis, respectively. Strains were grown in LB supplemented with NO_3_^-^ under anaerobic conditions for 24 hours, then switched to aerobic conditions with succinate as carbon source, i.e. in conditions wherein growth strictly relies on UQ (24). The wt, Δ*menA*, and the Δ*menA*Δ*ubiUV* strains showed differential efficiency in shifting from anaerobiosis to aerobiosis, the lag periods lasting 2 to 4 hours for the wt and Δ*menA* strains, and lasting 7 hours for the Δ*menA*Δ*ubiUV* mutant (Figure 8A). This indicated that UbiUV-synthesized UQ was important for allowing a fast transition, presumably as a consequence of a higher level of UQ in wt than in Δ*menA*Δ*ubiUV* mutant. Eventually, both strains showed the same growth rate in exponential phase and reached the same final OD_600_ value, suggesting that the UbiIHF-synthesized UQ was activated and fully compensated the requirement of UQ in extended aerobic conditions. To confirm this hypothesis, we re-inoculated these cells into the same medium (Figure 8A, refresh), and as expected we observed that lag periods were the same for both strains since they had accumulated the same level of UQ since the beginning of the growth. In contrast, the Δ*menAΔubiH* mutant – a strain defective for the aerobic UQ-synthesis pathway – exhibited drastic differences as compared with the two strains above, i.e. a slower and shorter exponential phase and a lower final OD_600_ value. Moreover, Δ*menA* Δ*ubiH* mutant failed to resume growth upon reinoculation in fresh medium (Figure 8A). Altogether these results indicated that the level of anaerobically UbiUV- synthesized UQ sustained the anaerobic-to-aerobic shift but failed to sustain protracted aerobic growth. This view was further supported by measuring the UQ content during transition from anaerobic to aerobic conditions in a separate experiment (Figure 8B). For this, cultures in LB of Δ*ubiUV* or Δ*ubiT* mutants were subjected or not to chloramphenicol (Clp) treatment prior to shift and samples were taken at 0 min, 30 min, and 120 min for UQ quantification. UQ level increased with time in both the wt and the Δ*ubiUV* mutant but in the 30-120 min period it stopped increasing in the presence of translation inhibitor Clp. The likeliest explanation is that UQ biosynthesis is driven by UbiUV before the shift and later de novo synthesized by UbiIHF in aerobic conditions. This suggested that the 3 hydroxylases UbiI, H, and F were already present under anaerobiosis, in a stand-by state, waiting for O_2_ to allow hydroxylation. Importantly, this was confirmed as levels of UbiI, H, and F proteins were found to be similar in both aerobic and anaerobic conditions (Figure S7).

**Figure 8.**
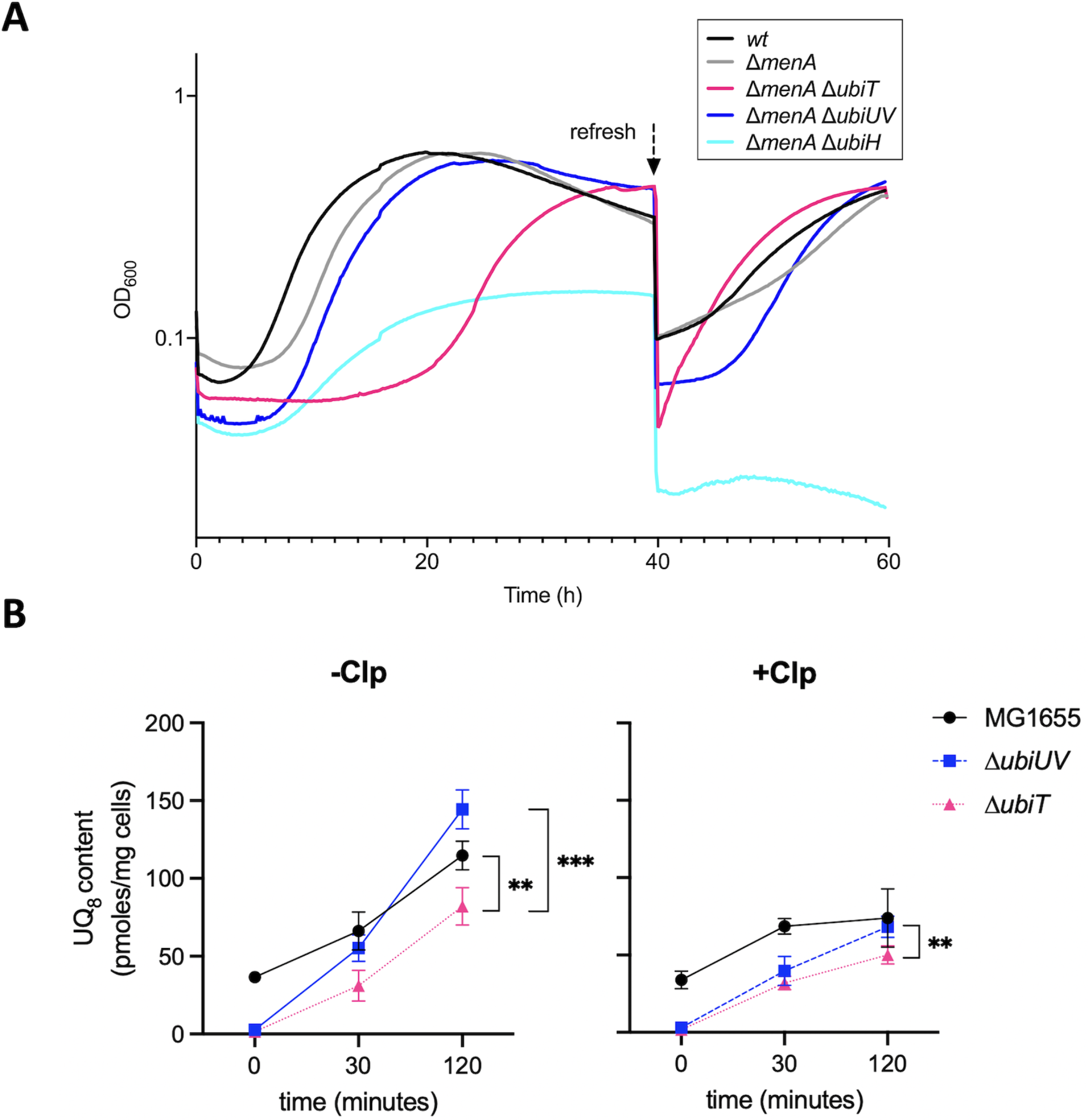
Role of *ubiUVT* in the anaerobic to aerobic transition. (A). *E. coli* wt and strains devoid of the MK/DMK *(ΔmenA)* and UQ aerobic (Δ*ubiH*) and anaerobic (Δ*ubiUV* or *ΔubiT*) synthesis pathways, were grown anaerobically in LB KNO_3_ medium, washed in M9 medium without carbon source and resuspended in M9 succinate medium to OD_600_=0.02. Growth was followed aerobically at 37°C in a TECAN microplate reader in 3 independent experiments. At 40 hours of growth, cells were diluted 1/100 in the same medium (refresh) and growth was resumed for 20h more. (**B).** *E. coli* wt (MG1655), Δ*ubiUV* and Δ*ubiT* strains were cultured anaerobically in LB medium containing NO_3_^-^ as final electron acceptor until OD ∼ 1. After 20 min of treatment with chloramphenicol (+Clp) at 200 µg/mL or without chloramphenicol (-Clp) under anaerobic conditions, the cultures were shifted to ambient air for a two-hour incubation. UQ_8_ content was quantified before (0 min) or after oxic transition (30 min and 120 min) by HPLC-ECD of lipid extracts from 1 mg of cells. Quantifications are expressed as picomole per milligram of cells (n=4 biological replicates). **, P < 0.01; ***, P < 0.001 by unpaired Student’s t test. Mean ± standard deviation is indicated.

Secondly, the role of the accessory factor, UbiT, was investigated using the Δ*menA* Δ*ubiT* mutant. As described before, the Δ*menA*Δ*ubiT* strain was grown first in LB with NO_3_ under anaerobiosis, subsequently shifted in succinate minimal medium, and growth was monitored. A most unexpected and spectacular effect was observed as lag period with this strain was 3 and 5 times longer than the one observed for the Δ*menA* Δ*ubiUV* and wt strains, respectively (Figure 8A). However, the Δ*menA*Δ*ubiT* strain finally reached a final OD_600_ value similar to WT, Δ*menA*, Δ*menA* Δ*ubiUV* strains at 40h, and also resumed growth upon re-inoculation at 40h (Figure 8A). This highlighted a crucial role of UbiT in the anaerobic-aerobic transition phase. This result was strengthened by direct quantification of UQ synthesized with time after shifting cultures from anaerobiosis to aerobiosis (Figure 8B). The Δ*ubiT* mutant exhibited a 2-fold reduction in UQ as compared with the Δ*ubiUV* mutant after the shift. When Clp was added, the difference was much smaller. This confirmed that UbiT is necessary at the onset of aerobic UQ biosynthesis, presumably via the UbiIHF complex.

### 8. The *yhbS* gene is not involved in UQ-based metabolism

The *yhbS* gene predicted to encode an acetyltransferase, lies downstream the *ubiT* gene (Figure S8A). It was recently proposed to intervene in sncRNA-mediated expression control (29). Using RT-PCR, we showed that *yhbS* and *ubiT* genes share a single transcription unit (Figure S8B). Using YhbS-SPA tag protein, we observed that YhbS protein synthesis takes place both under aerobiosis and anaerobiosis. The level of YhbS-SPA protein appears slightly higher in -O_2_ and this induction seems to be lost in the *Δfnr* mutant, as expected if *yhbS* and *ubiT* genes are co-expressed and co-regulated by Fnr (Figure S8C). The *ΔyhbS* mutant shows no defect in NO_3_^-^ respiratory capacity, and no aggravating effect was observed upon combining *ΔyhbS* and *ΔmenA* mutations (Figure S8D). Last, we carried out shift experiments, from -O_2_ to +O_2_, as described above for *ubiT* and failed to identify any defect in the *ΔyhbS* mutant (not shown). Altogether with previous assays failing to reveal a defect in UQ levels in anaerobiosis in the Δ*yhbS* mutant (3), these results allowed us to rule out a role of YhbS in UQ synthesis.

## CONCLUSION

UQ is an essential component of electron transfer chains, and of respiratory metabolism. For decades, the dogma has been that UQ was exclusively used for aerobic respiratory metabolism, whereas DMK/MK was used for electron transfer in anaerobic respiratory chains. Following our recent discovery that UQ is also synthesized under anaerobiosis, which contradicted the above assumption (3), the present study identified two versatile anaerobic physiological processes that rely on the anaerobic UQ biosynthesis pathway, namely NO_3_^-^ respiration and uracil biosynthesis. Moreover, we provide clear evidence that UbiUV catalyze hydroxylation steps independently from O_2_. Last, UbiT was found to play a key role in both anaerobiosis and aerobiosis conditions, allowing a smooth transition between the two conditions. Overall, this analysis uncovers a new facet of the strategy used by *E. coli* to adapt to changes in O_2_ level and respiratory conditions. This is of particular interest in the context of gut microbiota studies, as changes in O_2_ level and in respiratory electron acceptors are key factors that the host uses to select the type of flora present through the different sections of the intestine (30).

UbiUV-mediated UQ synthesis takes place under anaerobiosis. Here we showed that this is made possible by Fnr-mediated activation of expression of the *ubiUV* operon that takes place from microaerobiosis (0.1% O_2_) to anaerobiosis. In contrast, expression of the *ubiT* gene is more versatile with 2 promoters, one under Fnr control, allowing UbiT synthesis under micro- and an-aerobiosis, simultaneously with UbiUV, and the second constitutive one, insuring expression in aerobiosis. This genetic regulation is consistent with the presence of UbiT proteins under both aerobic and anaerobic conditions. Such a versatile expression meets with other evidence we collected, which together pave the way to an important role of UbiT in anaerobiosis to aerobiosis transition: (i) UbiT is required for insuring continuous UQ synthesis upon shifting from anaerobiosis to aerobiosis, (ii) *ubiT* was found to compensate for the lack of *ubiJ* in conditions where high dosage of *ubiUV* genes suppressed absence of *ubiIHF* under aerobiosis, (iii) UbiIHF enzymes are present in anaerobiosis but not active as one would expect for O_2_-dependent hydroxylases. This indicates that the O_2_- dependent pathway is in a stand-by mode in anaerobic conditions, waiting only for the presence of O_2_ to activate the O_2_-dependent hydroxylases and produce UQ, as proposed previously (31). This is also consistant with the fact that UbiUV synthesis is strictly controlled at the transcriptional level, whereas expression of *ubiIHF* is constitutive. Altogether, this leads us to propose that UbiT and UbiJ are required for the formation of two related but distinct metabolons, respectively an anaerobic one containing UbiUV, and an aerobic one containing UbiIHF. Besides, both UbiJ and UbiT are likely to bind UQ biosynthetic intermediates via their SCP2 domain, thereby providing the substrates to UbiUV and UbiIHF (9, 32).

UbiUV catalyze hydroxylation of the benzene ring in the absence of O_2_. Moreover, our results show that they can substitute to aerobic hydroxylases UbiIHF in the presence of O_2_, but that they still catalyze the hydroxylation without relying on O_2_ in this condition. This raises the question of the source of the O atom under anaerobiosis. Previous analysis on RhlA, a member of the U32 protein family to which UbiU and V belong, indicated that prephenate, an intermediate within the aromatic amino acid biosynthesis pathway, could act as O donor (11). Our ongoing studies aim at investigating such a possibility in the case of anaerobic UQ biosynthesis. [Fe-S] clusters seem to play a role in the process, since *isc* mutants devoid of anaerobic [Fe-S] biogenesis machinery and UbiU variant lacking [Fe-S] cluster fail to produce UQ. The simplest hypothesis is that [Fe-S] clusters are transferring electrons from the O source to a terminal reductase, both to be identified.

UbiUVT-synthesized UQ has a significant contribution to growth in anaerobiosis and in microaerobiosis (0.1% O_2_). Indeed, we found that UbiUVT-synthesized UQ are key for NO_3_^-^ respiration in the absence of (D)MK, in agreement with early biochemical work on formate-nitrate reductase (26) and with our previous study reporting that *Pseudomonas aeruginosa* denitrifying activity depends on UbiUVT synthesized UQ (32). Moreover, we observed that the anaerobically synthesized UQ greatly contributes to uracil synthesis. This was unexpected as uracil synthesis was reported to depend mainly on the oxidation of (S)- dihydroorotate to orotate with fumarate as hydrogen acceptor and DMK/MK as an electron carrier (26). Our present physiological studies demonstrate that the anaerobically produced UQ can fully compensate the DMK/MK loss, likely through an as yet unknown reductase since UQ is too electro-positive to be a FrdABCD substrate (33). Last, UQ could be used as an electron sink to other catabolic processes taking place in both aerobiosis and anaerobiosis such as heme biosynthesis, wherein HemG enzyme utilizes UQ or MK for the conversion of protoporphyrinogen IX into protoporphyrin IX (34).

The contribution of anaerobically synthesized UQ for *E. coli* multiplication in the gut appeared as marginal. This implies that either absence of UV-synthesized UQ was masked by MK/DMK synthesis, or anaerobic UQ-dependent processes such as NO_3_^-^ respiration or uracil biosynthesis is dispensable. Clearly the first possibility is the likeliest given the paramount importance of anaerobic respiration for *E. coli* multiplication in the gut (35, 36), as nicely confirmed by the drastically altered multiplication of MK/DMK deficient cells (Figure 7). This is of particular interest as presence and nature of respiratory electron acceptors were proposed to be drivers of bacterial community composition in the different regions of the intestine (30). Likewise, the relatively high O_2_ level in duodenum, of NO_3_^-^ in ilium, and hypoxia in cecum were proposed to be causal of the different flora hosted in these regions in healthy host. Strategies used by *E. coli* to live in such different respiratory and fermentative conditions are therefore key aspects of its adaptation to the host. In this context, it is important to understand the mechanism underlying the switch from O_2_-rich to NO_3_^-^-rich and/or hypoxic compartments and the present study highlights the added value of having overlapping systems permitting smooth shift from anaerobic NO_3_^-^ to aerobic respiration.

## MATERIAL AND METHODS

### Strain constructions (Table 1)

Most knockout strains were obtained by generalized Φ P1 transduction using donor strains from the Keio collection (37). For introducing the SPA-tag on the chromosome or for generation of specific knockouts, PCR recombination with the lambdaRed system was used, using the oligonucleotides indicated in Table 1 (18, 38). When necessary, the antibiotic resistance marker was removed using FLP recombinase expression from plasmid pCP20 as described previously (39). Cassette removal and plasmid loss were verified by antibiotic sensitivity and confirmed by PCR amplification. Point mutations were introduced on the chromosome using the pKO3 vector (40).

**Table 1:**
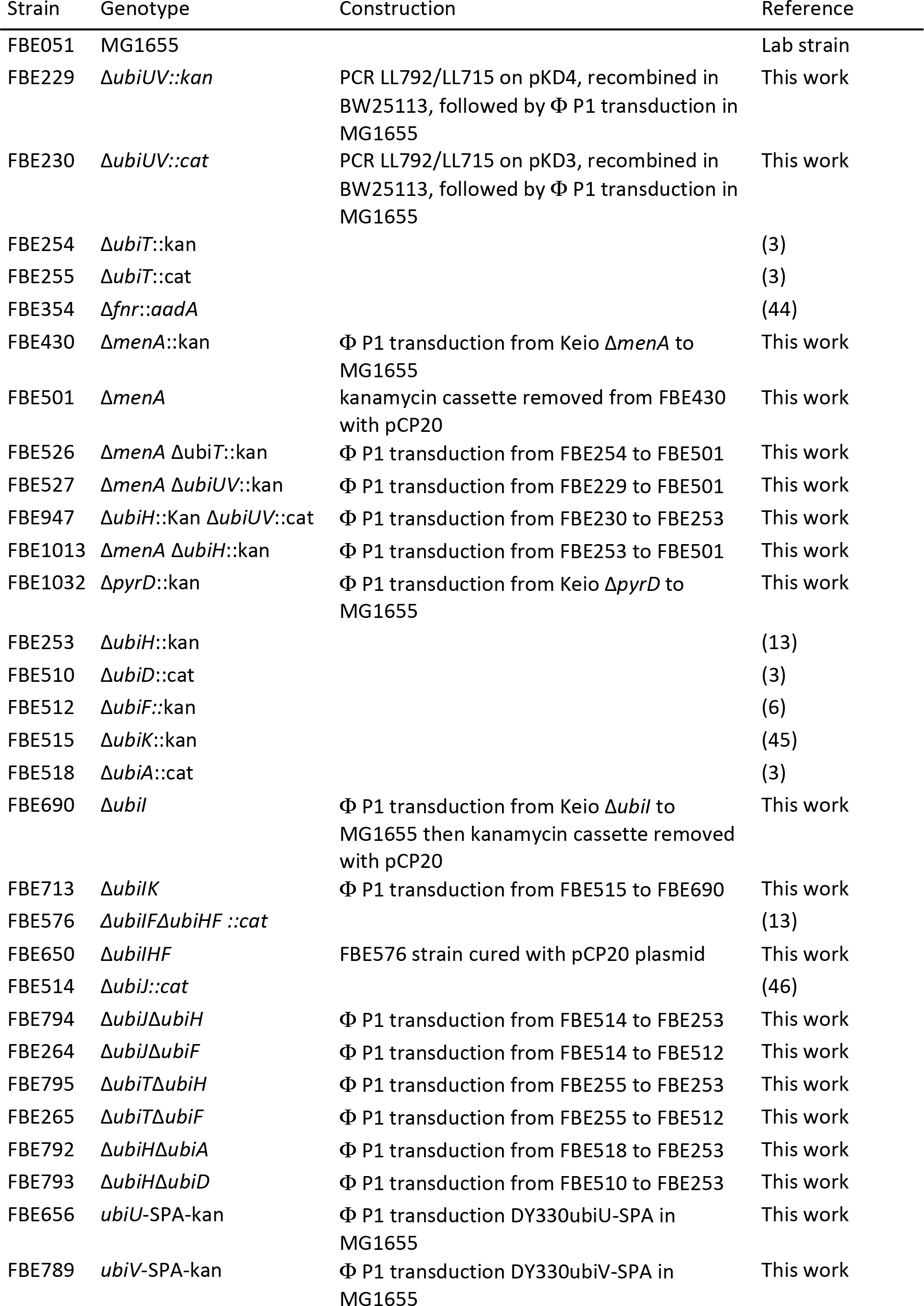
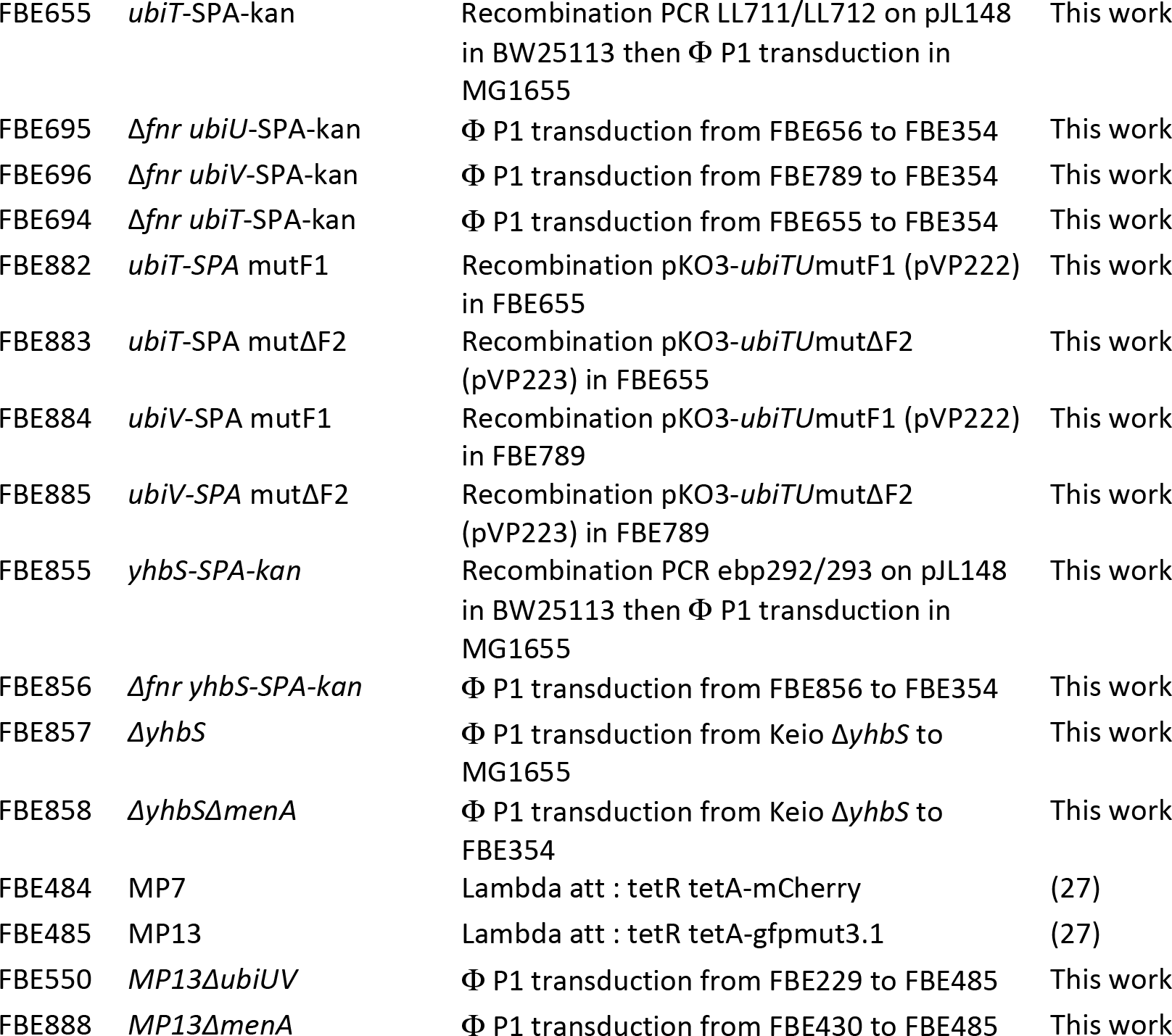
Strains used in this study

For mouse intestine colonization experiments, we used MP7 and MP13 strains, which derive from the commensal *E. coli* MP1 strain (27). MP7 and MP13 express respectively mCherry or GFP under the control of a tetracycline inducible promoter. Δ*menA* and Δ*ubiUV* deletions were introduced in MP13 using generalized Φ P1 transduction.

### Plasmid constructions (Table 2)

pUA66 and pUA-*ubiUV*p plasmids were obtained from the library of *E. coli* promoters fused to GFP coding sequence (20). The *ubiT* transcriptional fusions were constructed using primers indicated in Table 3 and cloned in XhoI/BamHI sites of pUA66. Expression plasmids for *ubiUV* and *fnr* were constructed using primers indicated in Table 3 and cloned in EcoRI/SalI sites of pBAD24 vector (41). Expression plasmids for *ubiIHF* and *ubiM_Neisseria* genes were constructed using primers indicated in Table 3 and cloned in EcoRI/XhoI sites of pTet vector. A region of 1275 base pairs encompassing *ubiU* and *ubiT* promoters was cloned in pKO3 vector (40). Mutations were introduced in the pKO3-*ubiTU* vector, in the pBAD- *ubiUV*, and in the transcriptional fusions by PCR mutagenesis on plasmid, using the oligonucleotides indicated (Tables 2 and 3).

**Table 2:** Plasmids used in this study.

| Plasmid | Name | Description / Construction | Source |
| --- | --- | --- | --- |
| pCP20 | pCP20 | Ap, Cm, FLP recombinase expression | (39) |
| pEB227 | pBAD24 | AmpR – ColE1 ori – PBAD promoter | (41) |
| pEB267 | pKD46 | AmpR – ts ori – lambda Red genes | (38) |
| pEB268 | pKD3 | AmpR – FRT-cat-FRT | (38) |
| pEB269 | pKD4 | AmpR – FRT-kanaR-FRT | (38) |
| pEB794 | pJL148 | AmpR – SPA-FRT-kanaR-FRT | (18) |
| pES154 | pBAD-*ubiUV* | PCR on MG1655 genomic DNA *ebp134/136* (EcoRI/XhoI) in pBAD24 (EcoRI/SalI) | This work |
| pES185 | pBAD-*ubiU*(C176A)*V* | Mutagenesis PCR *ebp178/179* on pES154 | This work |
| pES184 | pBAD-*ubiUV*-SPA | PCR *ebp134/ebm968* on strain FBE696(EcoRI/XhoI) in pBAD24 (EcoRI/SalI) | This work |
| pVP040 | pBAD-*fnr* | PCR on MG1655 genomic DNA *ebp31/32* (MfeI/XhoI) in pBAD24 (EcoRI/SalI) | This work |
| pEB1242 | pASK-IBA37plus | AmpR – ColE1 ori – TetR promoter – 6His | IBA |
| pEB1823 | pTet | PCR mutagenesis *ebm1567/1568* on pEB1242 to remove 6His tag | This work |
| pES060 | pTet-*ubil* | PCR on MG1655 genomic DNA *ebp64/65* (EcoRI/XhoI) in pTet (EcoRI/XhoI) | This work |
| pES059 | pTet-*ubiH* | PCR on MG1655 genomic DNA *ebp61/62* (EcoRI/XhoI) in pTet (EcoRI/XhoI) | This work |
| pES143 | pTet-*ubiF* | PCR on MG1655 genomic DNA *ebp37/38* (EcoRI/XhoI) in pTet (EcoRI/XhoI) | This work |
| pES151 | pTet-*ubiM*_Neisseria | PCR on MG1655 genomic DNA *ebp139/140* (EcoRI/XhoI) in pTet (EcoRI/XhoI) | This work |
| pEB898 | pUA66 | KanR - pSC101 ori - GFPmut2 | (20) |
|  | pUA-*ubiUVp* |  | (20) |
| pVP220 | pUA-*ubiUVp*mutF1 | Mutagenesis PCR *Ebp287/288* on pUA-*ubiU* | This work |
|  | pUA-*ubiTp1p2* |  | (20) |
| pVP169 | pUA-*ubiTp1* | PCR on MG1655 genomic DNA *ebp191/192* (XhoI/BamHI) in pUA66 (XhoI/BamHI) | This work |
| pVP170 | pUA-*ubiTp2* | PCR on MG1655 genomic DNA *ebp193/194* (XhoI/BamHI) in pUA66 (XhoI/BamHI) | This work |
| pVP187 | pUA-*ubiTp1p2ΔF2* | Mutagenesis PCR *Ebp237/238* on pUA-*ubiT* | This work |
| pVP221 | pUA-*ubiTp1p2mutF2* | Mutagenesis PCR *Ebp289/290* on pUA-*ubiT* | This work |
| pEB232 | pKO3 | camR, pSC101 ori, *sacB* | (40) |
| pVP219 | pKO3-*ubiTU* | PCR on MG1655 genomic DNA *ebp276/291* (XhoI/BamHI) in pKO3 (SalI/BamHI) | This work |
| pVP222 | pKO3-*ubiTU*mutF1 | Mutagenesis PCR *ebp287/288* on pVP219 | This work |
| pVP223 | pKO3-*ubiTU*mutΔF2 | Mutagenesis PCR *ebp237/238* on pVP219 | This work |

**Table 3:**
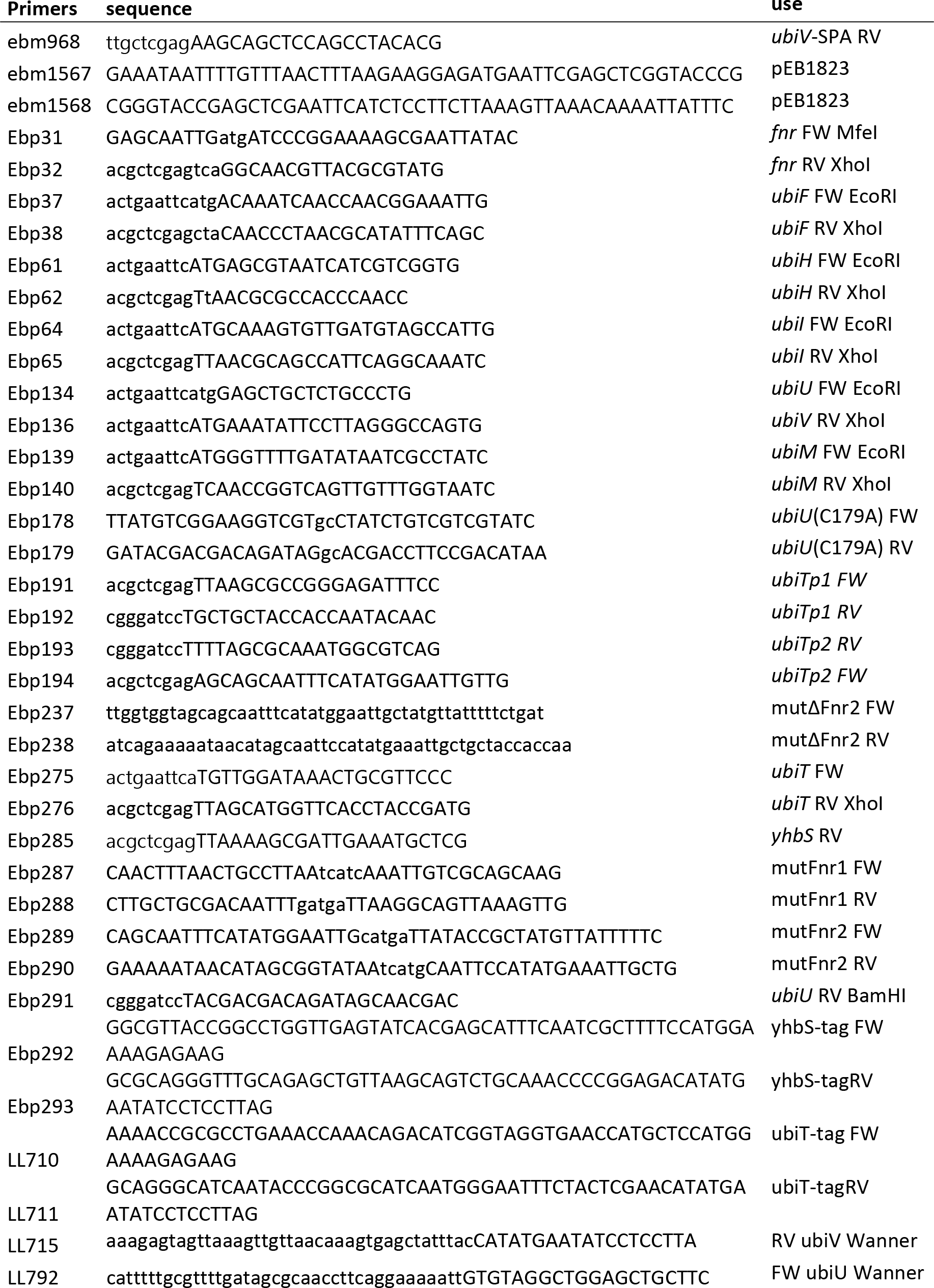
Primers used in this study.

### Media and growth conditions

Strains were grown in LB miller (10g/l of tryptone, 10g/l of NaCl and 5g/l of yeast extract) or M9 medium (6 g/l Na_2_HPO_4_•7H_2_O, 3 g/l KH_2_PO_4_, 0.5 g/l NaCl, 1 g NH_4_Cl, 2 mM MgSO_4_, 1mg/ml thiamine) supplemented with 0.2% glucose, 0.2% glycerol, or 50mM succinate as the carbon source. For anaerobic cultures, media were degazed and incubated in anaerobic environment for at least 24 hours prior to use, if necessary supplemented with 25mM KNO_3_ as electron acceptor and uracil 25ug/ml or casamino acids at 0.05%.

For microaerobic experiments, media and plates were pre-equilibrated and cells were cultured in a Whitley® H35 hypoxic station with 95% N2, 5% CO_2_ and the desired O_2_ concentration. Humidity and temperature were set up at 85% and 37°C, respectively. For anaerobic-aerobic shift experiments, all anaerobic steps were performed in a JACOMEX® Campus anaerobic chamber under N_2_ atmosphere at 1ppm O_2_ maximum. Cells were first isolated anaerobically in LB agar plates supplemented with 0.2% Glucose and incubated overnight at 37°C. Next day, cells were cultured anaerobically in 3mL LB supplemented with 25mM NO3^-^ for 24 hours at 37°C. Still under anaerobiosis, cells were collected by centrifugation, supernatant was discarded, and pellets were washed twice using 1mL M9 medium without carbon source and normalized at 0.1 OD units in M9 medium supplemented with 50mM Sodium Succinate. At this point, cultures were moved out to atmospheric air and growth was followed by triplicate at 37°C on 200µl of culture in a 96- well plate using a TECAN infinite M200 plate reader. At 40h of culture, cells were diluted 1/20 in new M9 50mM Sodium Succinate medium and readings were resumed until 60 hours.

Aerobic and anaerobic cultures for quinone analysis:

For aerobic cultures, 5 mL of LB medium, supplemented with Ampicillin (100 µg/mL) and 0.05% arabinose when necessary to induce the expression from the pBAD vectors, was inoculated with 100 µl of an overnight culture in glass tubes (15 cm long and 2 cm in diameter) and incubated at 37°C, 180 rpm overnight.

Anaerobic cultures were performed in Hungate tubes as previously described (3). Briefly, LB medium was supplemented with 100 mM KNO_3_ as final electron acceptor, 100 mg/liter L- cysteine (adjusted to pH 6 with NaOH) in order to reduce residual molecular oxygen and 2.5 mg/liter reasazurin. This medium was distributed in Hungate tubes and deoxygenated by high purity argon bubbling for 40 min. The Hungate tubes were sealed and autoclaved. The resazurin was initially purple, it turned to pink after deoxygenation and become colorless after autoclave. The preculture was performed overnight at 37°C in Eppendorf tubes filled to the top with LB medium containing 100 mM KNO_3_. The Hungate tubes were then inoculated through the septum by disposable syringes and needles with 100 µL of precultures and incubated at 37°C without agitation. The resazurin remained colorless during culture indicating anaerobic conditions.

For anaerobic to aerobic shift assay, MG1655 WT, Δ*ubiUV*, and Δ*ubiT* strains were grown anaerobically in Hungate tubes for ∼ 4 hours. Then, 26 µL of chloramphenicol (200 µg/mL) was injected through the septum by Hamilton syringe. After 20 minutes, the Hungate tubes were unsealed, 2 mL of cultures was taken for lipid extraction and quinone analysis. The rest of cultures was transferred to 250 mL Erlenmeyer flasks and placed at 37°C, 180 rpm for 2 hours. 2 mL aliquots of cultures were taken at 30 min and 120 min after transition to ambient air for lipid extraction and quinone analysis.

### SDS-PAGE and Western blotting

Total cell extracts were prepared by resuspending cell pellets in Laemli buffer 1X at a concentration of 0.3 OD_600nm_ units in 10 µl, and then heating for 10 minutes at 95°C. After separation of 8 µl of total cell extracts on SDS-PAGE, electrotransfer onto nitrocellulose membranes was performed using Trans-Blot turbo transfer system from Biorad. After blocking in PBS 1X + milk 5%, SPA-tagged proteins were detected with monoclonal anti-Flag M2 antibody purchased from Sigma. YbgF protein was used as an internal control and revealed with polyclonal anti-YbgF antibodies. Fluorescent secondary antibodies were respectively IRDye 800 anti-mouse and IRDye 680 anti-rabbit purchased from Li-Cor. Scanning and quantification were performed on a Li-Cor Odyssey-Fc imaging system, reading at 700 nm (for YbgF detection) or 800 nm (for Flag detection).

### Transcriptional fusions with GFP

We used several clones from the *E. coli* transcriptional fusions library (20) and we constructed the required additional transcriptional fusions (see above for plasmid construction and Table 2). Δ*fnr E. coli* strain was co-transformed with plasmids carrying the *gfp* transcriptional fusions and compatible pBAD24 or pBAD-*fnr* plasmids. Selection plates were incubated at 37°C for 16h. 600 µl of LB medium supplemented with kanamycin and ampicillin, and with 0.02% arabinose for pBAD-driven expression, were incubated (4 biological replicates for each assay) and grown for 16 hours at 37°C in 96-well polypropylene plates of 2.2 ml wells in anaerobiosis. Cells were pelleted and resuspended in PBS supplemented with 30 µg/ml chloramphenicol and incubated at 4°C for 1 hour before fluorescent intensity measurement was performed in a TECAN infinite M200 plate reader. 150 µl of each well was transferred into black Greiner 96-well plate for reading optical density at 600nm and fluorescence (excitation: 485nm; emission: 530 nm). The expression levels were calculated by dividing the intensity of fluorescence by the optical density at 600 nm, after subtracting the values of a blank sample. These results are given in arbitrary units because the intensity of fluorescence is acquired with an automatic optimal gain and hence varies from one experiment to the other.

### Lipid extraction and quinone analysis

Cultures of 2, 5, or 10 mL were cooled on ice for at least 30 min before centrifugation at 3200 x g at 4°C for 10 min. Cell pellets were washed in 1 mL ice-cold phosphate-buffer saline (PBS) and transferred to preweighted 1.5 mL Eppendorf tubes. After centrifugation at 12,000 g at 4 °C for 1 min, the supernatant was discarded, the cell wet weight was determined and pellets were stored at -20°C until lipid extraction, if necessary. Quinone extraction from cell pellets was performed as previously described (6). The dried lipid extracts were resuspended in 100 µL ethanol, and a volume corresponding to 1 mg of cell wet weight was analyzed by HPLC electrochemical detection-MS (ECD-MS) with a BetaBasic- 18 column at a flow rate of 1 mL/min with a mobile phase composed of 50% methanol, 40% ethanol, and 10% of a mix (90% isopropanol, 10% ammonium acetate (1 M), and 0.1% formic acid). When necessary, MS detection was performed on an MSQ spectrometer (Thermo Scientific) with electrospray ionization in positive mode (probe temperature, 400°C; cone voltage, 80 V). Single-ion monitoring detected the following compounds: UQ_8_ (M+H^+^), m/z 727-728, 6–10 min, scan time of 0.2 s; 3(^18^O)-UQ_8_ (M+H^+^), m/z 733-734, 6–10 min, scan time of 0.2 s; UQ_8_ (M+NH_4_^+^), m/z 744-745, 6–10 min, scan time of 0.2 s; UQ_10_ (M+NH_4_^+^), m/z 880– 881, 10-17 min. MS spectra were recorded between m/z 600 and 900 with a scan time of 0.3 s. ECD and MS peak areas were corrected for sample loss during extraction on the basis of the recovery of the UQ_10_ internal standard and then were normalized to cell wet weight. The peaks of UQ_8_ obtained with electrochemical detection or MS detection were quantified with a standard curve of UQ_10_ as previously described (6).

### 18O2 labeling

MG1655 wt and Δ*ubiI*Δ*ubiH*Δ*ubiF* containing respectively the pBAD24 empty vector or pBAD-*ubiUV* were grown overnight at 37°C in LB medium supplemented with Ampicillin (100 µg/mL) and 0.05% arabinose. These precultures were used to inoculate 20 mL of the same fresh medium at an optical density at 600 nm (OD_600_) of 0.05 in Erlenmeyer flasks of 250 mL. The cultures were grown at 37°C, 180 rpm, until an OD_600_ of 0.4-0.5 was reached. An aliquot was taken for lipid extraction and quinone analysis (0 min of ^18^O_2_) and 13 mL of each culture was transferred to an Hungate tube. 5 mL of labeled molecular oxygen (^18^O_2_) was injected through the septum with disposable syringes and needles, and the incubation was continued at 37°C, 180 rpm for 2 hours. Then 5 mL of each sample was taken for quinone analysis (120 min of ^18^O_2_).

### Mouse intestine colonization experiments

All animal experiments were performed in accordance with the institutional and national guidelines. Experiments were performed under the supervision of C.L. (agreement 38 10 38) in the Plateforme de Haute Technologie Animale (PHTA) animal care facility (agreement C3851610006 delivered by the Direction Départementale de la Protection des Populations) and were approved by the ethics committee of the PHTA and by the French government (APAFIS#14895- 2018042623275606.v5).

4-week-old female BALB/cByJ were purchased from Charles River Laboratories (Saint-Germain-Nuelles) and were acclimatized in a controlled animal facility under specific pathogen-free conditions for two weeks prior to the beginning of the colonization assay. Mice were randomly assigned to groups of three or five per cage and ear punching was used in order to identify each mouse in a given cage.

The colonization experiments were adapted and performed as previously described (42, 43). Mice were given drinking water containing streptomycin sulfate and glucose (both 5 g/L) for 72 hours to remove existing resident anaerobic facultative microflora. For clearance of streptomycin, fresh water devoid of antibiotic and glucose was then given to mice for 48 hours before inoculation of *E. coli* strains and for the rest of experiment. To start the competition experiment, the mice were orally inoculated with 200 µL of a mixture in a 1:1 ratio of the two competing strains at ∼ 20,000 cells/mL in PBS. Mice from each cage were orally inoculated with the same solution of bacteria. An aliquot of inoculum was plated on LB agar containing 15 µg/mL tetracycline in order to compute the input value.

The relative abundance of both competing strains was then monitored at several days post-inoculation in fecal samples. Fecal samples were collected from each mouse in preweighed 1.5 mL Eppendorf tubes containing the equivalent of 100 µL glass beads (diameter 0.25 to 0.5 mm) and 80 µL PBS and the feces weight was determined. A volume of PBS was then added to each tube in order to obtain a final concentration of 0.15 g of feces per 1 mL PBS. The feces were homogenized by vortexing for 2 min, serially diluted by 10-fold steps up to a 10^5^-fold dilution, and aliquots of 70 µL were plated on LB agar medium containing 15 µg/mL tetracycline. The plates were incubated overnight at 37°C and were transferred at 4°C for at least 2 hours the following day, before imaging under blue light which revealed the fluorescent markers carried by each colony. The red and green colonies corresponding respectively to MP7 and MP13 strains were counted by an adapted version of ImageJ. Then, the CFU was computed per gram of feces for each strain and a competitive index (CI) was calculated as a ratio of (MP13 mutant CFU/MP7 wt CFU) / (input MP13 mutant CFU/input MP7 wt CFU), where the input CFU was determined from the inoculum for which an aliquot was plated on the day of gavage. The limit of detection in fecal plate counts was 10^2^ CFU/g feces. At all-time points, the wt strain was detectable on the fecal plates. The absence of CFU count and CI for one day in one mouse corresponds to the absence of feces for that day. Significance of CI was calculated by GraphPad Prism using one-sample t test compared to one.

## Acknowledgements

We thank Marc Fontecave and Murielle Lombard from College de France, and the members of the SAMe unit at Pasteur for discussion and help. We thank Mark Goulian (University of Pennsylvania, USA) for providing the MP7 and MP13 *E. coli* strains and Laurent Loiseau for providing the UbiUVT SPA-tagged strains. We gratefully acknowledge the help of TrEE team members with the mouse intestine colonization experiments, Françoise Blanquet, Dalil Hannani, Clément Caffaratti, and Amélie Amblard. We are also grateful to Arnold Fertin for developing the ImageJ plugin used for the automatic counting of red and green colonies. This project was supported by Institut Pasteur and CNRS and by grants from the ANR (ANR- 10-LABX-62-IBEID and ANR-19-CE44-0014O2-TABOO).

---

Figure S1. **O_2_-dependent and O_2_-independent biosynthetic pathways of UQ in *E. coli*.** R, octaprenyl chain illustrated on the UQ_8_ structure; 4-HB, 4-hydroxybenzoic acid; OPP, 3- octaprenylphenol; DMQ_8_, C-6-demethoxyubiquinone; DMAPP, dimethylallyl pyrophosphate; IPP, isopentenyl pyrophosphate. The Ubi-enzymes and accessory factors common between the two pathways are in black, those corresponding to the aerobic pathway in red, and those corresponding to the anaerobic pathway are in blue. Hydroxyl groups added on C5, C1, and C6 are highlighted in red.

Figure S2. Amount of UbiV-SPA produced in +O_2_ from pBAD-UbiUV-SPA plasmid compared with physiological amount in –O_2_ of chromosome-encoded UbiV-SPA. Strain UbiV-SPA (FBE789) was grown in LB in the absence of O_2_. Wild type *E. coli* transformed by pBAD- UbiUV-SPA (pES184) was grown in LB in +O_2_ and induced 2 hours with 0.02% arabinose. After preparation of whole cell extract, the sample was diluted 2-fold serially (1 to 1/32). Western blot was performed using an anti-Flag antibody. ni : uninduced cells.

Figure S3. Ubiquinone 8 (UQ_8_), demethylmenaquinone 8 (DMK_8_) and menaquinone 8 (MK_8_) content in the indicated mutant strains of the Keio collection (37) after anaerobic growth overnight at 37°C in LB medium. Mean ± standard deviations (SD) (n=2).

Figure S4. **Comparison of UbiT-SPA levels in the regulation mutants in –O_2_ and in +O_2_.** *E. coli* strains expressing UbiT-SPA with or without the Δ*fnr* or mutΔF2 chromosomal mutations (FBE655, FBE694, and FBE883) were grown in biological duplicates in LB at 37°C in the indicated oxygenic conditions until OD_600nm_=1. Normalized quantities of total protein extracts in duplicate were separated by SDS-PAGE 12% and detected by Western-Blot using anti-Flag monoclonal antibody for the detection of the SPA tag or anti-YbgF polyclonal antibodies as an internal loading control.

Figure S5. **(A)** Ubiquinone 8 (UQ_8_), **(B)** demethylmenaquinone 8 (DMK_8_) and **(C)** menaquinone 8 (MK_8_) content in MP7 wt, MP13 Δ*ubiUV,* MP13 Δ*menA* strains after anaerobic **(A)** and aerobic **(B and C)** growth overnight at 37°C in LB medium. Mean ± standard deviations (SD) (n=2). N.D., not detected.

Figure S6. Total CFU count per gram of feces (**A and B**) and competitive index (**C and D**) for either MP7 (mCherry-tagged MP1) WT:MP13 (GFP-tagged MP1) Δ*ubiUV* (**A and C**) or MP7 WT:MP13 Δ*menA* (**B and D**) competition experiments in each mouse of the experiments shown in Figure 7. The limit of detection was 10^2^ CFU. The absence of total CFU count in one day corresponds to the absence of feces for that day.

Figure S7. **Protein levels of UbiIHF proteins in aerobic and anaerobic conditions.** Strains producing UbiI, UbiH, and UbiF tagged with SPA at the chromosome, were grown in LB in aerobic and anaerobic conditions. Whole cell extracts were analyzed by Western blot with an anti-Flag antibody. Results representative of two independent experiments.

Figure S8. ***ubiT* is in operon with the unknown function *yhbS* gene. (A).** Genetic organization. See legend of figure 5. (**B).** RT-PCRs were performed on total RNA prepared on MG1655 cells in exponential phase, with oligonucleotides ebp275 and ebp285 (Table 3). The positions of hybridization of the oligonucleotides are indicated in panel A. +/- RT indicates the absence or presence of the reverse transcriptase (RT) enzyme in the reaction mixture. A control PCR was performed on genomic DNA with the same oligonucleotides. **(C).** *E. coli* strains YhbS-SPA and Δ*fnr*/YhbS-SPA (FBE855, FBE856) were grown in LB at 37°C in the indicated oxygenic conditions until OD_600nm_=1. Normalized quantities of total protein extracts in duplicate were separated by SDS-PAGE 12% and detected by Western-Blot using anti-Flag monoclonal antibody for the detection of the SPA tag. **D.** The indicated *E. coli* strains were grown anaerobically for two days at 37°C on M9 medium plates supplemented with 0.2% glycerol and NO_3_.

## Notes

### Competing Interest Statement

The authors have declared no competing interest.

