## Supplementary figures S1 to S8 for "Role of the *Escherichia coli* ubiquinone-synthesizing UbiUVT pathway in adaptation to changing respiratory conditions"

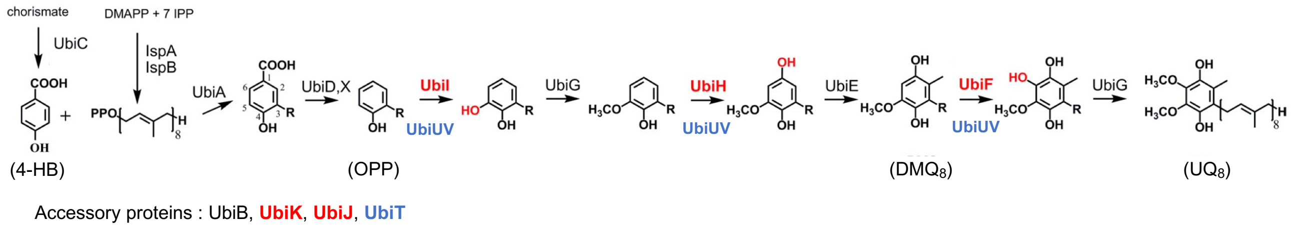


**Figure S1 : $O_2$-dependent and $O_2$-independent biosynthetic pathways of UQ in *E. coli*.** R, octaprenyl chain illustrated on the $UQ_8$ structure; 4-HB, 4-hydroxybenzoic acid; OPP, 3-octaprenylphenol; $DMQ_8$, C-6-demethoxyubiquinone; DMAPP, dimethylallyl pyrophosphate; IPP, isopentenyl pyrophosphate. The Ubi-enzymes and accessory factors common between the two pathways are in black, those corresponding to the aerobic pathway in red, and those corresponding to the anaerobic pathway are in blue. Hydroxyl groups added on C5, C1, and C6 are highlighted in red.

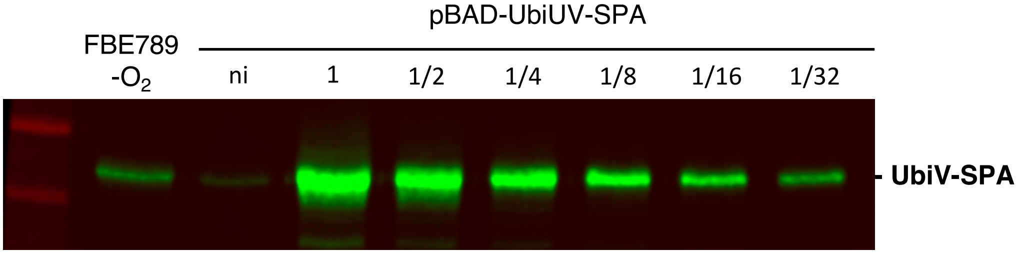


**Figure S2 : Amount of UbiV-SPA produced in $+O_2$ from pBAD-UbiUV-SPA plasmid compared with physiological amount in $–O_2$ of chromosome-encoded UbiV-SPA.** Strain UbiV-SPA (FBE789) was grown in LB in the absence of $O_2$. Wild type *E. coli* transformed by pBAD-UbiUV-SPA (pES184) was grown in LB in $+O_2$ and induced 2 hours with 0.02% arabinose. After preparation of whole cell extract, the sample was diluted 2-fold serially (1 to 1/32). Western blot was performed using an anti-Flag antibody. ni : uninduced cells.

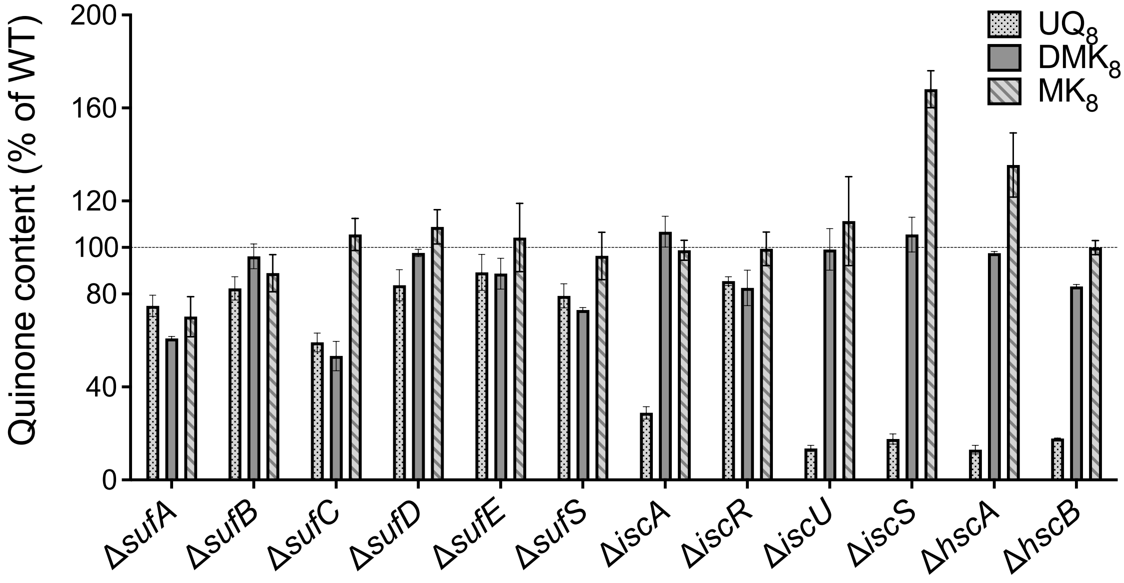


**Figure S3 :** Ubiquinone 8 ($UQ_8$), demethylmenaquinone 8 ($DMK_8$) and menaquinone 8 ($MK_8$) content in the indicated mutant strains of the Keio collection (37) after anaerobic growth overnight at 37°C in LB medium. Mean ± standard deviations (SD) (n=2).

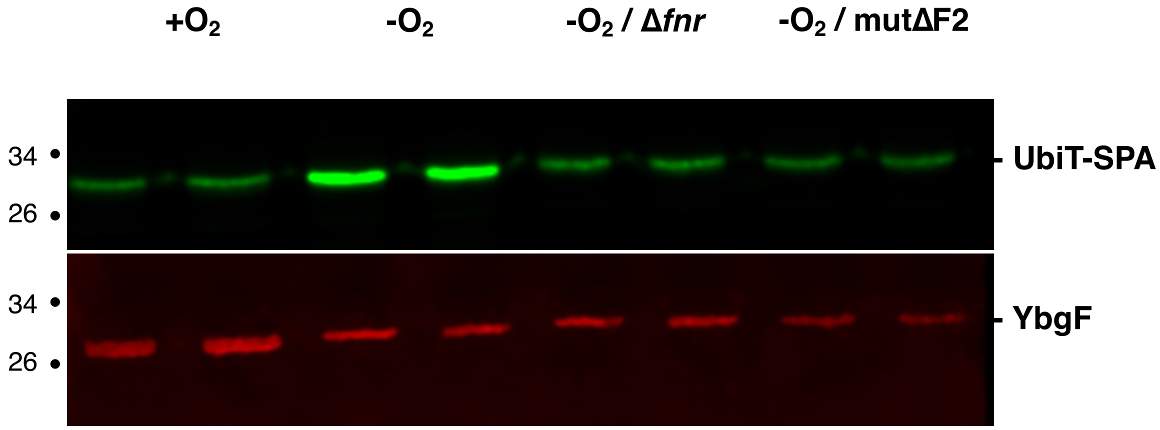


**Figure S4 : Comparison of UbiT-SPA levels in the regulation mutants in –$O_2$ and in +$O_2$.** *E. coli* strains expressing UbiT-SPA with or without the Δ*fnr* or mutΔF2 chromosomal mutations (FBE655, FBE694, and FBE883) were grown in biological duplicates in LB at 37°C in the indicated oxygenic conditions until $OD_{600nm}$=1. Normalized quantities of total protein extracts in duplicate were separated by SDS-PAGE 12% and detected by Western-Blot using anti-Flag monoclonal antibody for the detection of the SPA tag or anti-YbgF polyclonal antibodies as an internal loading control.

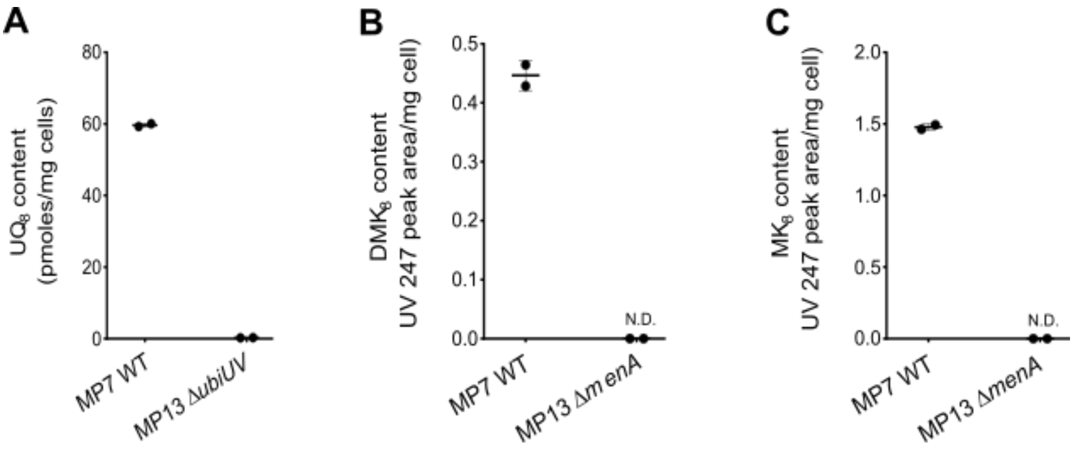


**Figure S5:** (**A**) Ubiquinone 8 ($UQ_8$), (**B**) demethylmenaquinone 8 ($DMK_8$) and (**C**) menaquinone 8 ($MK_8$) content in MP7 WT, and MP13Δ*ubiUV* or MP13Δ*menA* strains after anaerobic (**A**) and aerobic (**B and C**) growth overnight at 37°C in LB medium. Mean ± standard deviations (SD) (n=2). N.D., not detected.

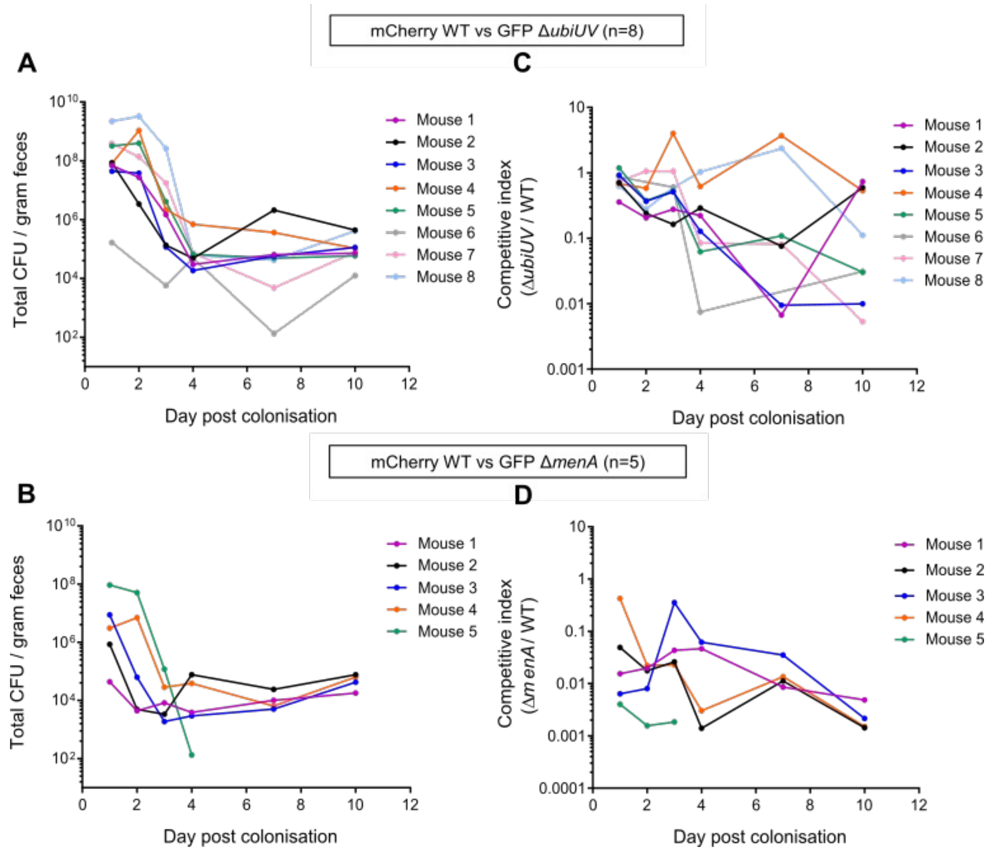


**Figure S6:** Total CFU count per gram of feces (**A and B**) and competitive index (**C and D**) for either MP7 (mCherry-tagged MP1) WT:MP13 (GFP-tagged MP1) Δ*ubiUV* (**A and C**) or MP7 WT:MP13 Δ*menA* (**B and D**) competition experiments in each mouse of the experiments shown in Figure 7. The limit of detection was $10^2$ CFU. The absence of total CFU count in one day corresponds to the absence of feces for that day.

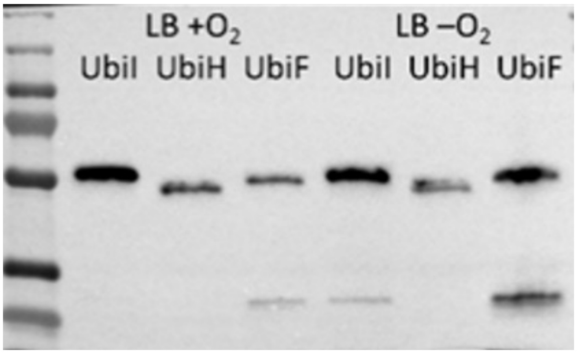


**Figure S7: Protein levels of UbiIHF proteins in aerobic and anaerobic conditions.** Strains producing UbiI, UbiH, and UbiF tagged with SPA at the chromosome, were grown in LB in aerobic and anaerobic conditions. Whole cell extracts were analyzed by Western blot with an anti-Flag antibody. Results representative of two independent experiments.

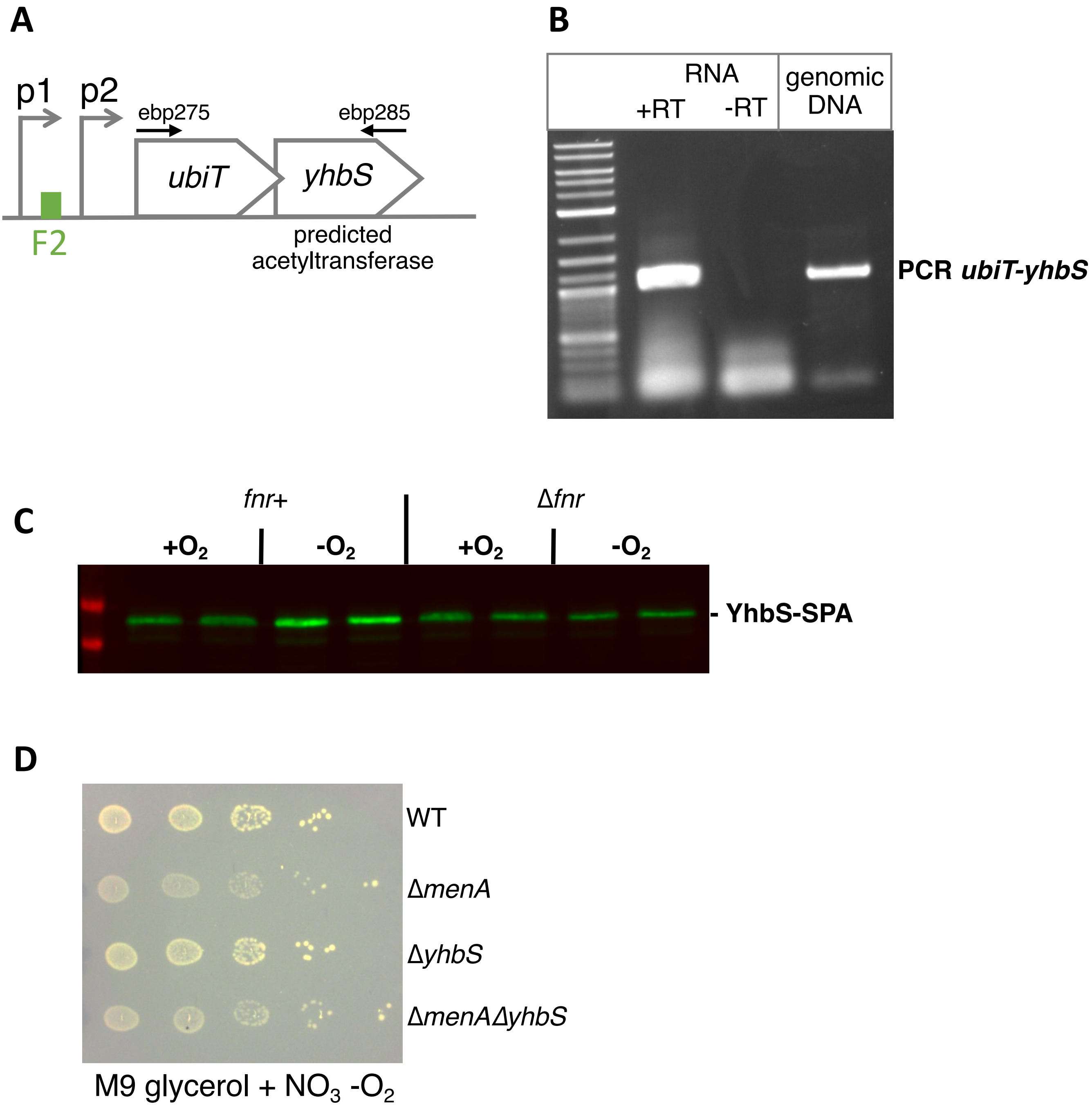


**Figure S8 : *ubiT* is in operon with the unkown function *yhbS* gene. A.** Genetic organization. See legend of figure 5. **B.** RT-PCRs were performed on total RNA prepared on MG1655 cells in exponential phase, with oligonucleotides ebp275 and ebp285 (Table 3). The positions of hybridization of the oligonucleotides are indicated in panel A. +/- RT indicates the absence or presence of the reverse transcriptase (RT) enzyme in the reaction mixture. A control PCR was performed on genomic DNA with the same oligonucleotides. **C.** *E. coli* strains YhbS-SPA and Δ*fnr*/YhbS-SPA (FBE855, FBE856) were grown in LB at 37°C in the indicated oxygenic conditions until $OD_{600nm}$=1. Normalized quantities of total protein extracts in duplicate were separated by SDS-PAGE 12% and detected by Western-Blot using anti-Flag monoclonal antibody for the detection of the SPA tag. **D.** The indicated *E. coli* strains were grown anaerobically for two days at 37°C on M9 medium plates supplemented with 0.2% glycerol and $NO_3$.
